# Structural analysis reveals high-order Nipah Virus Fusion Protein assemblies and interactions with neutralising nanobodies

**DOI:** 10.64898/2026.05.22.727168

**Authors:** Joseph W. Thrush, Luke Jones, Irene Carlon-Andres, Miles Carroll, Sergi Padilla-Parra, Raymond J. Owens

## Abstract

Nipah virus (NiV) is a highly pathogenic emerging henipavirus for which there are no licensed vaccines or therapeutics currently available. Viral entry is mediated by the coordinated action of the attachment (NiVG) and fusion (NiVF) glycoproteins yet the organisation of NiVF on viral particles and its influence on antibody neutralisation is not completely understood. Using cryo-electron tomography of NiV pseudovirions and virus-like particles (VLPs), we directly observe inter-F contacts on viral membranes. These interactions occur frequently but are distributed stochastically, forming variably sized assemblies rather than ordered arrays, consistent with transient or low-affinity associations. We have generated single domain antibodies (termed nanobodies) to the NiVF glycoprotein that bind to an epitope at the interface between neighbouring NiVF trimers and therefore occlude the inter-NiVF contacts. We show that these nanobodies inhibit VLP cell fusion and neutralise NiV pseudoviruses *in vitro*, which suggests that the interaction between NiVF trimers is important for NiV virus entry. The identification of the inter-NiVF contacts on viral particles adds a previously unrecognised structural feature to the organisation of the NiV entry machinery which could be targeted to prevent viral infection.

## Main

Nipah virus (NiV) is a highly pathogenic henipavirus in the *paramyxoviridae* family, first discovered in a patient presenting with encephalitis in 1998 in Malaysia^1,2^. Infected individuals present with dizziness, fatigue, vomiting and diarrhoea, which often develops into severe encephalitis or respiratory conditions, resulting in a high mortality rate (40-70%) in symptomatic individuals^2–4^. The natural reservoir is *Pteropus* fruit bats, which are distributed over south-east Asia, the Indian sub-continent and eastern Africa^4,5^. Transmission occurs through consumption of contaminated date palm sap, or with contact with infected pigs^4^. Analysis of NiV outbreaks has revealed two major genetic lineages, Malaysian and Bangladesh, which have a 91.8% sequence homology, occurring in south-east Asia and the Indian sub-continent respectively^6,7^. Other henipavirus cases, however, have occurred elsewhere, such as Hendra virus in Australia^8^ and the recently discovered Langya virus in China^9^.

NiV is a negative-sense single-stranded RNA virus, with nine encoded proteins, two of which, the Glycoprotein (NiVG) and the Fusion (NiVF) protein, are responsible for host-receptor binding and host-viral membrane fusion respectively^10,11^. NiVF is a class I fusion protein, synthesised as a F_0_ precursor and is cleaved by cathepsin L into F_1_ and F_2_ subunits in the producer cell^12^. Mature NiVF exists as a metastable prefusion homotrimer, similar to other *paramyxoviridae* fusion proteins. NiVG is a type II transmembrane protein that binds host receptor EphrinB2^11^. Receptor binding of NiVG causes triggering of NiVF, which undergoes a cascade of conformational changes through the prefusion, intermediate state, and culminating in the collapse of the heptad repeat bundle into the post-fusion conformation and fusion of the viral and host membranes, which releases virion contents into the cell^13,14^.

The mechanism of NiV infection, in particular the protein interactions on the viral membrane responsible for host-cell membrane fusion, is still unclear^13,14^. Two competing models have been posited to describe the interactions between NiVF and NiVG: the *clamp* model and the *provocateur* model^14–19^. The *clamp* model postulates that NiVG interacts with NiVF before receptor binding, effectively stabilising the pre-fusion conformation. The *provocateur* model postulates that NiVG interacts with NiVF after receptor binding, stimulating NiVF to transition from the pre-fusion to the intermediate conformation.

The determination of the structure of NiVF by either X-ray crystallography^20^ or cryo-electron microscopy (cryo-EM) structure^21^ suggests the presence of NiVF-NiVF interactions on the surface of NiV. The crystal structure shows that NiVF forms a hexamer-of-trimers with C3 symmetry and two different interfaces within the hexamer. In the cryo-EM structure, NiVF-NiVF also exists as a dimer-of-trimers; however, the dimer interface is not the same as in the hexamer in the crystal structure but corresponds to the interface between adjacent hexamers observed by crystal packing. Therefore, the two papers disagree on which NiVF-NiVF interface may be biologically relevant, and therefore which could be exploited for therapeutic gain by preventing or disrupting NiVF dimer formation.

NiVF is an important immunodominant target for neutralising antibodies and is highly conserved between NiV genotypes and the related Hendra virus, making it an attractive candidate for vaccine design^6^. The majority of neutralising agents against NiVF are monoclonal antibodies^22–27^; however, nanobodies or VHHs (Variable Heavy-chain domains of Heavy-chain only antibodies) offer a viable alternative as antiviral agents with several advantages compared to conventional antibodies^28,29^. Nanobodies are smaller in size, significantly more stable than whole antibodies and can be produced in bacterial systems, reducing reagent costs. Nanobodies are single-domain binding proteins but can also be expressed recombinantly fused to IgG Fc domains to generate bivalent constructs with increased functional affinity due to a gain in avidity. Recently, neutralising nanobodies against NiVF have been reported that bind to a quaternary epitope crypt on the head region of the protein^30^. By coupling the most potent nanobody, DS90, to an anti-NiVG neutralising antibody^31^ a bispecific reagent was produced which was anticipated to have greater resistance to viral escape than anti-NiVF or NiVG antibodies alone^30^. Focusing on NiVF, we have combined structural analysis and nanobody generation to expand on the mechanism of NiV viral entry, in particular the protein interactions involving NiVF on the viral membrane using NiV viral-like particles and Lentiviral pseudoviruses. Using cryo-tomography, we have generated the first full-length NiVF trimeric structure resolved as a dimer-of-trimers embedded within a NiV VLP and provide evidence that NiVF trimers form higher order assemblies in the VLP membrane. We have produced neutralising nanobodies targeting the NiVF glycoprotein that map to the NiVF dimer interface, suggesting a new mechanism for blocking infection through disrupting NiVF assemblies in the viral membrane.

## Results

### Generating Nipah Virus Model Systems

A NiV pseudovirus consisting of plasmids expressing full-length NiVF and NiVG (Malaysian genotype), alongside a HIV-1 GagPol plasmid, p8.91, and a firefly luciferase reporter plasmid, CSFLW was used as a model for authentic NiV, which can only be handled at BSL4. A NiV VLP model was also established by co-expression in HEK293T cells of the full-length NiVF and NiVG plasmids (Malaysian genotype), together with a NiV Matrix (NiVM) plasmid generated in house. NiVM was recombinantly fused with a beta-lactamase enzyme, which acted as the reporter protein for VLP-cell fusion^32^. Both models were used for structural analysis by cryo-ET and in nanobody characterisation assays.

### Cryo-Electron Tomography of NiV Pseudoviruses

In order to understand the structural traits of NiVF and NiVG, which drive NiV fusion, we employed cryo-electron tomography (cryo-ET) of NiV pseudoviruses decorated with both NiVF and NiVG Briefly, NiV pseudoviruses were generated, concentrated, applied to cryo-EM grids, plunge frozen, and tilt series collected at selected locations using a 300 kV cryo-electron microscope. Reconstructed tomograms showed NiV pseudoviruses exhibiting the typical HIV-1 core; mature virions exhibited cleaved GagPol forming a matrix protein lattice and an oblong capsid (Extended Data Fig. 1a-b). NiVF and NiVG decorated the pseudoviruses with high density, with over 100 envelope proteins counted on larger pseudoviruses. Sub-tomogram averaging of NiVF revealed a sub-population of NiVF in a dimer-of-trimer conformation, resolved to a resolution of 24.2 Å with C1 symmetry (Extended Data Fig. 1c).

### Cryo-Electron Tomography of NiV VLPs

To confirm whether the NiVF dimer-of-trimers was independent of HIV-1 Gag, cryo-ET of NiV VLPs comprising NiVF, G and M was carried out. The ratio of NiVF and NiVG plasmids combined with a fixed amount of NiVM for VLP formation was optimised to maximise fusogenicity of VLPs; NiVF being much more abundant than NiVG (Extended Data Fig. 2a). Reconstructed tomograms revealed a highly pleomorphic enveloped VLP, with highly dense decorations of NiVF and NiVG. Tomograms were collected of NiV VLP with sizes varying from 28 to 468 nm and these were non-uniform in shape and size, with variable sphericity (Fig. 1a-c). VLP thickness was limited to approximately 100 nm; likely flattened during the blotting and freezing process to accommodate them in the thin ice. The incorporation of NiVF and NiVG appears to be very stochastic, with vesicles containing either a majority NiVF, majority NiVG or a mixture of the two proteins. No NiVM lattices were observed in the VLPs which may be due to size constraints, or the lattices not forming due to the presence of the beta-lactamase reporter (BlaM)^32^.

**Figure 1:**
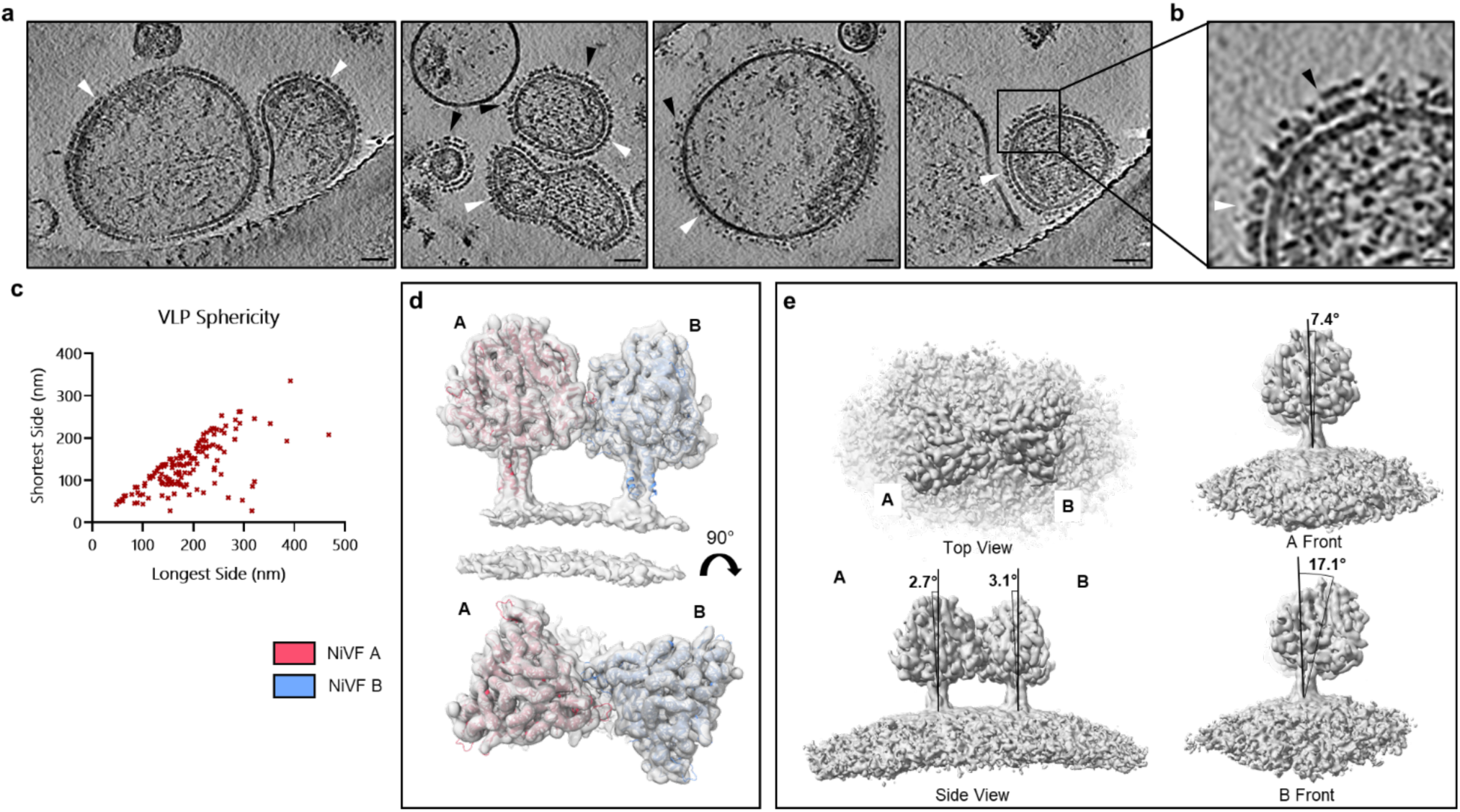
Cryo-ET of VLP and NiVF Dimer. (a) NiV VLP micrograph images showing pleomorphic nature of VLP with differences in both size and morphology. Incorporation of glycoproteins appears to be stochastic with VLPs varying from majority F to majority G. NiVF and NiVG are labelled with a white and black arrows respectively. Scale bar is 50 nm. (b) Zoomed in image of NiVF and NiVG on VLP. Scale bar is 15 nm. (c) VLP sphericity graph with longest side plotted against shortest side. Some VLPs are spherical while others are elongated and bulbous. (d) Sub-tomogram averaged volume of NiVF dimer-of-trimers with atomic model of Cryo-EM map fitted. (e) NiV dimer-of-trimers volume with C1 symmetry revealing asymmetricity. One of the dimers (A) is relatively normal to the membrane and the other is angled further towards the membrane (B). They are twisted 25° degrees relative to each other.

Sub-tomogram averaging of NiVF revealed a NiVF-NiVF dimer-of-trimers that occurred on the surface of the VLPs; resolved to a resolution of 7.2 Å with C1 symmetry (Fig. 1c). NiVFs in the dimer-of-trimers were relatively straight with minimal bending towards or away from each other, however there was significant twisting, approximately 25°, with one of the NiVF significantly more bent relative to the membrane at approximately 17°. The ability of NiVF to bend may be attributable to residue Pro486, which borders the ectodomain and the transmembrane domain, and disrupts the alpha-helix of a full-length AlphaFold2 model^33^(Extended Data Fig. 2e).

In previous work published work, the self-association of NiVF has been observed in X-ray crystallography or cryo-electron microscopy structures of both stabilised and non-stabilised isolated NiVF ectodomains structures^20,21^, however this is the first structure in which membrane bound NiVF dimers have been detected. There are three interfaces in the crystal structure of NiVF, two within a hexamer of NiVFs, and the third between neighbouring hexamers. The published cryo-EM structure of the NiVF dimer showed the interface in solution corresponding to the third interface between adjacent hexamers in crystal packing. The NiVF dimer observed here by cryo-ET of VLPs most closely matches the dimer in the cryo-EM structure, but is slightly more open at the apical end, which could be attributed to curvature forces imposed by the membrane (Fig. 2a). The alpha helices distal from the membrane are further apart in the cryo-ET volume than in the cryo-EM structure, with shifts of around 6.9 Å for a loop and 5.4 Å for a central alpha helix (Fig. 2b-e). The density around the cathepsin L cleavage site and the beginning of the fusion peptide was not resolved, similar to cryo-EM structures of isolated ectodomains^21,24–27^, suggesting that this region is still flexible in the dimer. This also means it is unclear whether NiVF is in the F_0_ precursor form, has been cleaved by cathepsin L into F_1_ and F_2_ subunits, or is a mixture of the two forms. There was unaccounted for density basal to the Cathepsin L cleavage site, possibly where the residues downstream of the cleavage site are located (Extended Data Fig. 2f).

**Figure 2:**
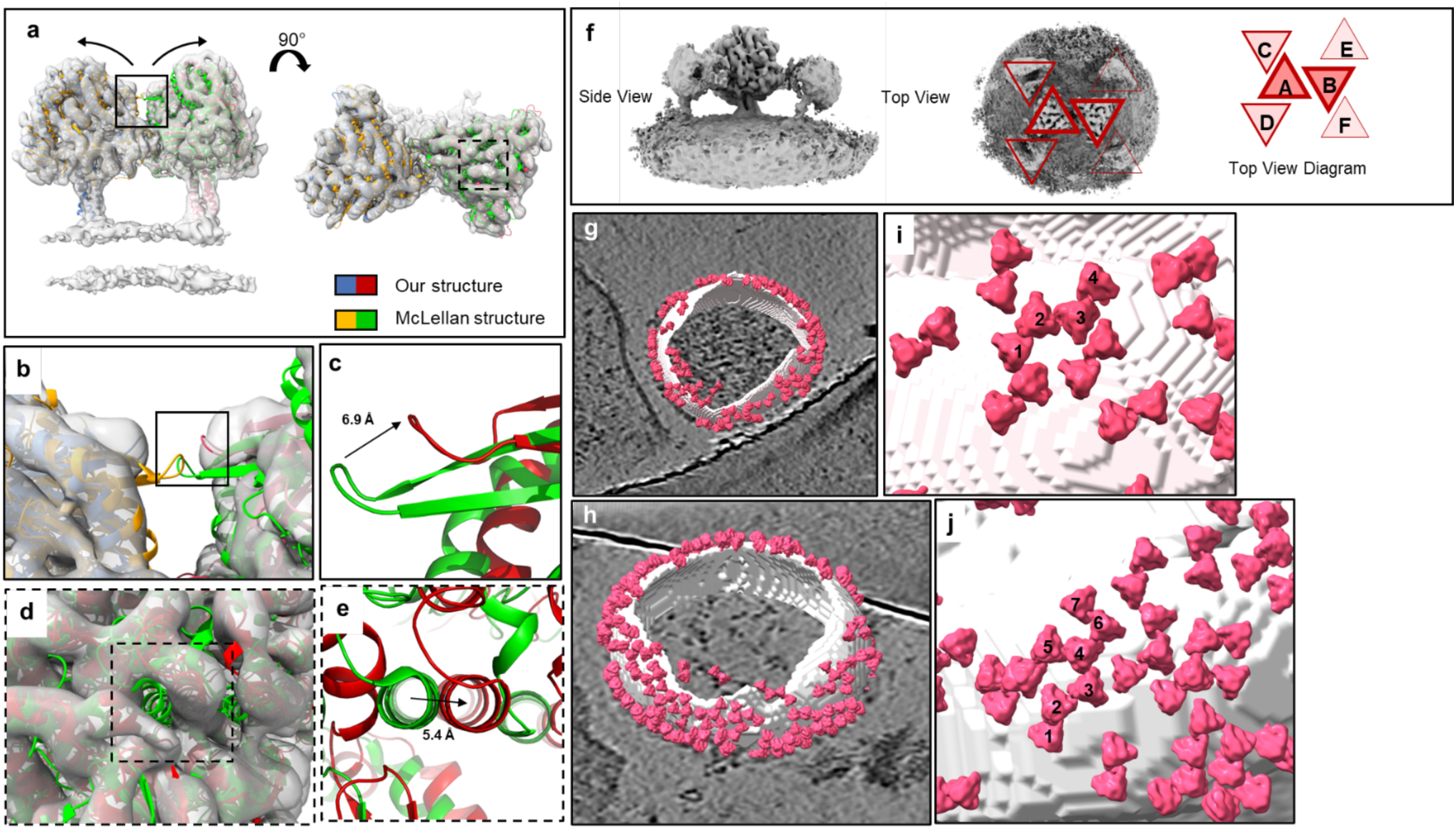
NiVF-Dimer in the context of the VLP. (a) Comparison of published cryo-EM structure^21^ with the cryo-ET STA volume. The cryo-ET volume reveals the NiVF trimers rotated away from each at the apex compared to the purified protein structure. The cryo-ET structure is in blue and red, and the cryo-EM structure is in yellow and green. (b-c) A shift of 6.9 Å of a looped region between the two structures. (d-e) A shift of 5.4 Å of an alpha helix between the two structures. (f) A diagram illustrating the NiVF network, with A and B forming the NiVF dimer-of-trimers in the cryo-ET volume, and lower density C,D,E,F being present in a sub-population of the sub-tomograms. There is a higher density next to NiVF A, suggesting it is more perpendicular because it needs to be tilted relative to those NiVF trimers (C and D). (g-h) Superimposing the NiVF Dimer subtomograms onto a VLP with the segmented membrane. Not all NiVF are labelled due to removal during classification, which has removed NiVF trimers or dimers-of-trimers that poorly matched the average. (i-j) Close up view of NiVF dimer-of-trimer subtomograms showing the extended nature of the interactions. There is a NiVF tetramer-of-trimers, labelled 1-4, with adjacent NiVF dimer-of-trimers and a heptamer-of-trimers labelled 1-7.

The asymmetrical twisting of the NiVF dimer-of-trimers is likely due to the NiVF sub-tomogram averaged structure not being a true dimer, but extending out to a trimer, tetramer or longer oligomer. By increasing the volume threshold of the structure, it was possible to see nearby NiVF at low-densities, suggesting the dimer-of-trimers does not occur in isolation, but rather an oligomeric network across the VLP surface (Fig. 2f). Superimposing the NiVF dimer-of-trimers sub-tomograms onto the VLP surface, the network was observed to extend to higher order oligomers, likely surrounded by NiV dimer-of-trimers and unbound, free NiVF trimers (Fig. 2g-j).

### Isolation and characterisation of NiVF specific nanobodies

To interrogate the NiVF dimer interface, antibodies to the pre-fusion conformation of NiVF (Malaysian genotype) were raised in a llama by three rounds of intra-muscular immunisation with purified pre-fusion stabilised NiVF ectodomain. NiVF ectodomain was stabilised in the pre-fusion conformation by the introduction of a disulfide bond, a rotation-restricting proline, a space-filling phenylalanine and a C-terminal GCNt trimerization domain, as previously described^34^. Peripheral blood mononuclear cells (PBMCs) were isolated, a phage display library VHH library was generated from the cDNA and subjected to two rounds of bio-panning against the stabilised NiVF ectodomain. The enrichment for VHHs that bound to NiVF was confirmed by a polyclonal phage ELISA and 93 phage clones from each round of panning were selected at random and tested for NiVF binding by ELISA. 26 clones from pan 1 and 60 from pan 2 with the strongest ELISA signals were sequenced which resulted in 27 clusters and 13 singlets, based on unique CDR3 sequences. 42 clones were expressed in *E. coli* and ranked based on their signal in a pre-fusion NiVF anti-VHH ELISA. 12 clones with high signal to noise in the ELISA were cloned into a recombinant-Fc vector, expressed in expi293™ and purified using immobilized metal affinity chromatography (IMAC). Binding of all 12 purified nanobody-Fcs to NiVF was confirmed by ELISA (Extended Data Fig. 3a-c).

Epitope binning of the 12 selected clones using biolayer interferometry (BLI) revealed two distinct epitopes, with a 10:2 split in nanobody binders indicating an immunodominant epitope on the NiVF (Extended Data Fig. 3d,e). Two nanobodies were selected from the epitope of the majority of the identified nanobodies (D2 and F10), and one from the minor epitope (G10) for further characterisation (Fig. 3). The binding affinities of monomer and bivalent-Fc constructs of D2, F10 and G10 to pre-fusion NiVF were measured by BLI and all three bound with low-to-sub nanomolar affinity (Fig. 4d and Extended Data Fig. 4b). D2 has a comparable K_D_ to the other published anti-NiVF nanobody (SPR K_D_ of 4.83 nM^30^, F10 has a 7-fold improvement and G10 has a comparable K_D_. Increased affinities were observed for the bivalent Fc fusions due to avidity gain: particularly D2 and G10 (Extended Fig. 4c).

**Figure 3:**
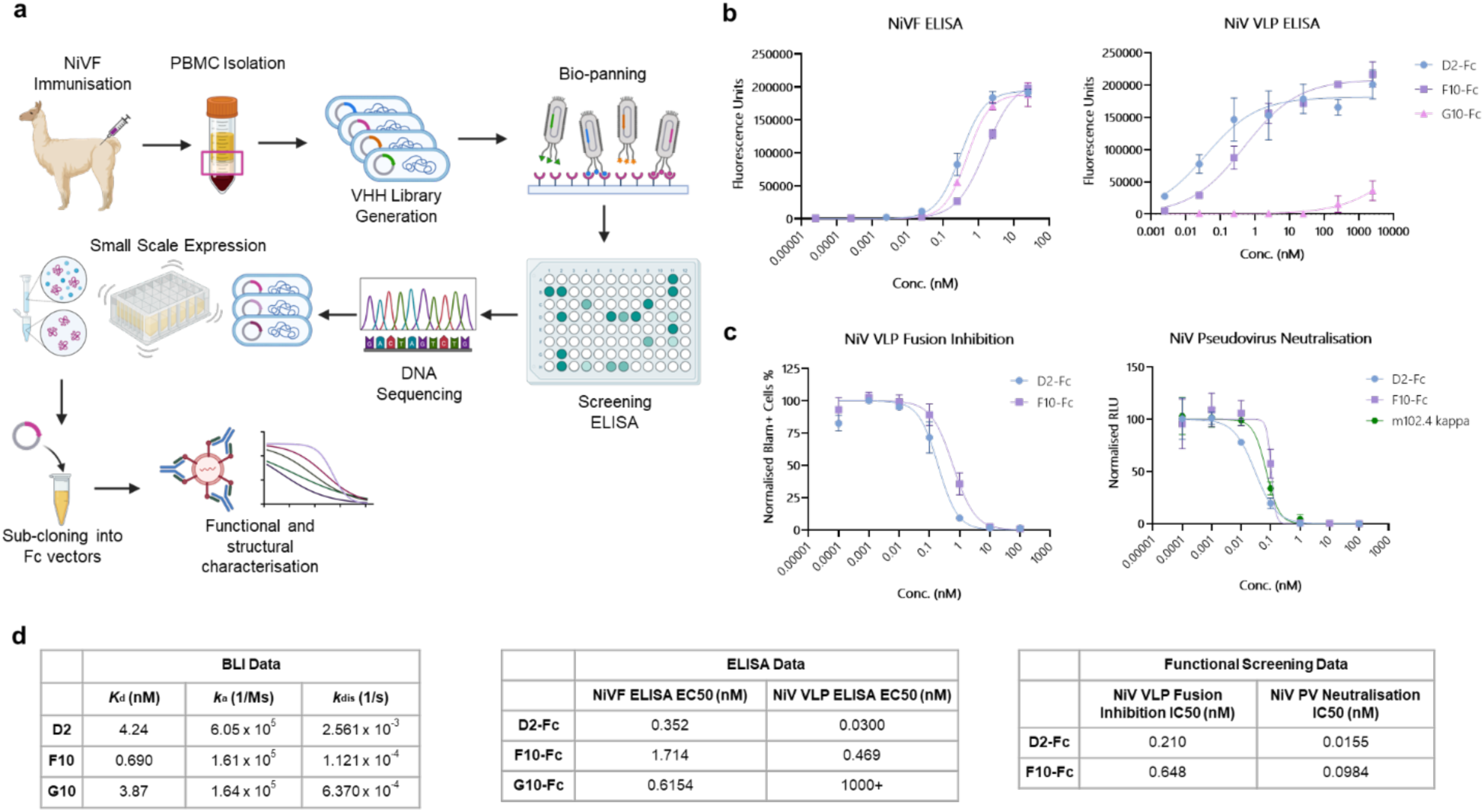
Nanobody discovery pipeline and characterisation. (a) VHH isolation and characterisation workflow. Immunisation of a llama with prefusion-stabilised NiVF ectodomain, followed by creation of a VHH library. Phage display is used to enrich the NiVF-specific VHH which are then screened with an anti-M13 ELISA. The positive hits are clustered by CDR3 after DNA sequencing and expressed as monomers in a small-scale bacterial culture and again screened with an anti-VHH ELISA. The positive hits are cloned into recombinant-Fc vectors and expressed in a mammalian expression system. They are screened for neutralisation and fusion inhibition activities using HIV-1 pseudovirus and NiV VLP model systems. Image made in BioRender. (b) ELISAs of Nb-Fc constructs against purified NiVF ectodomain and NiV VLP. Graph illustrates one of two repeats. (c) Fusion inhibition dilution curve of NiV VLP using D2-Fc and F10-Fc purified proteins. Neutralisation curves of NiV pseudovirus using D2-Fc and F10-Fc purified proteins. (d) Statistics table of binding affinity (Kd), association-rate (ka) and dissociation-rate (kdis) as calculated using the Octet Analysis Studio (v12.2.2.26). Summary of EC50 values of Nb-Fc constructs in ELISA assays. Values are calculated as the mean of two experiments, each with three repeats. Statistics table of VLP fusion inhibition and pseudovirus neutralisation assays. IC50 were calculated using two experiments, each with three technical repeats.

**Figure 4:**
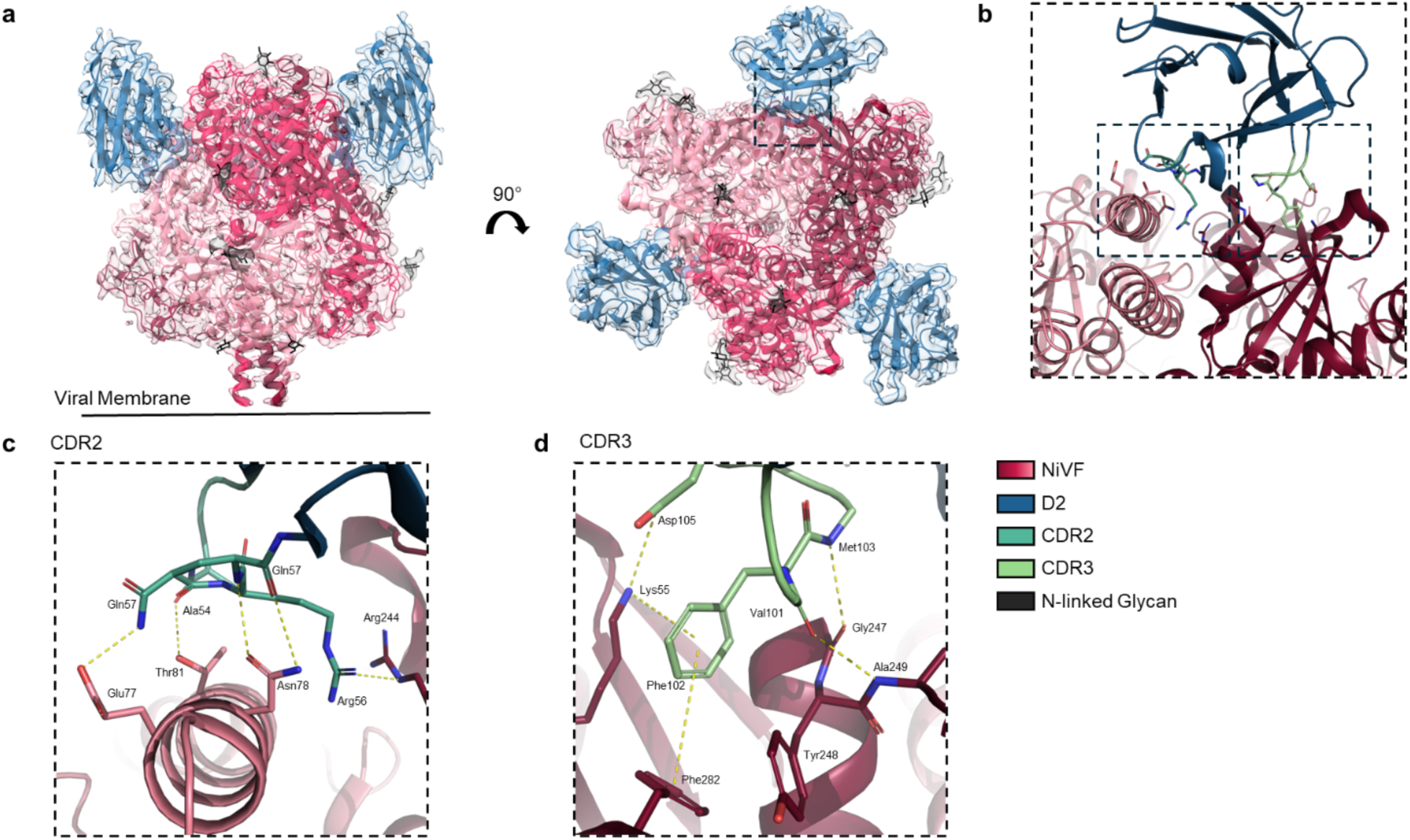
Epitope of D2 Nanobody Complex. (a) Model fitting of NiVF and D2 into the Cryo-EM SPA volume, NiVF in red, D2 in blue and glycans in black. (b) Zoomed in view of the NiVF-D2 epitope. (c) Close up view of the CDR2 of D2 interacting with an alpha helix and Arg of NiVF. (d) Close up view of the CDR3 of D2 fitting into a hydrophobic pocket of NiVF. NiVF is in shades of pink, D2 is blue and the CRDs are in shades of green. Side chains of interacting residues are shown as sticks with hydrogen bonds and pi interactions labelled with dashes.

We compared the binding in a NiVF ectodomain ELISA and a NiV VLP ELISA (Fig. 3b), as well as live-cell imaging and fluorescence lifetime cross-correlation spectroscopy (FLCCS)^35,36^. FLCCS is a single molecule resolution technique and therefore has increases sensitivity compared to ELISAs. FLCCS confirmed the ELISA results, by showing a high cross-correlation and a large slow-diffusion coefficient of D2 and F10 when mixed with an EGFP-labelled NiV pseudovirus suggesting D2 and F10 bound to NiVF on cells, VLPs and pseudovirions. Similarly to BLI, monomeric F10 bound with higher affinity than D2 to NiVF on pseudoviruses, suggesting the K_D_ values from pre-fusion stabilised NiVF accurately translate to native NiVF. A lack of cross-correlation of G10 suggests it only bound to the stabilised NiVF ectodomain (Extended Fig. 5,6). We conclude that the epitope recognised by G10 is either not present or is inaccessible on the VLP or pseudovirus.

### Fusion inhibition and neutralisation using a NiV VLP and HIV-1 pseudovirus model systems

As G10 did not bind to the VLPs it was not tested in the VLP fusion assay which showed that D2-Fc and F10-Fc inhibited fusion of the NiV Malaysian VLPs with IC50 values of 0.210 nM and 0.648 nM respectively (Fig. 3c,d). NiV HIV-1 pseudovirus infection of HEK293T cells was used to measure neutralisation capabilities in which the nanobodies showed similar IC50 values of 0.0155 nM and 0.0984 nM, respectively, comparable to a commercial anti-NiVG antibody, m102.4 kappa, run as a positive control in the assay^31^ (Fig. 3c,d). The similar potencies of the nanobodies in both VLP fusion and pseudovirus neutralisation assays suggests that the mechanism of inhibition involves the prevention of virus-cell fusion. The two virus models differ in the matrix proteins incorporated into the VLPs (NiVM) and pseudoviruses (HIV-1 Gag). The fact that comparable results were obtained in the two assays, showed that the fusion mechanism is not dependent on the source of the matrix protein.

### Structural analysis of NiVF-Nb complexes

Single particle analysis using cryo-electron microscopy (cryo-EM) was used to determine where the three nanobodies (D2, F10 and G10) bound to the ectodomain of stabilised NiVF. The nanobodies and NiVF protein were mixed in two different combinations, NiVF + D2 and NiVF + F10 + G10, vitrified on cryo-EM grids, and micrographs were collected for analysis. The NiVF-D2 and NiVF-F10-G10 complexes were resolved to resolutions of 2.59 Å and 2.95 Å respectively with C3 symmetry. Both D2 and F10 were observed bound to a quaternary epitope at the interface of two adjacent NiVF trimers (Fig. 4,5). Binding of a NiV neutralising nanobody, DS90 at this location has previously been reported^30^ but details of the pose and interacting residues differ from both D2 and F10. By contrast, G10 bound in the region of a loop (residues 103-106) containing the disulphide bridge (Cys104-Cys114) introduced by the mutations Leu104Cys and Ile114Cys to stabilise the NiVF ectodomain trimer^34^ and the Cathepsin L cleavage site, which is uncleaved in the purified protein (Extended Data Fig. 7,8). This would explain why G10 bound to the stabilised NiVF trimer but not the native NiVF on the VLPs.

**Figure 5:**
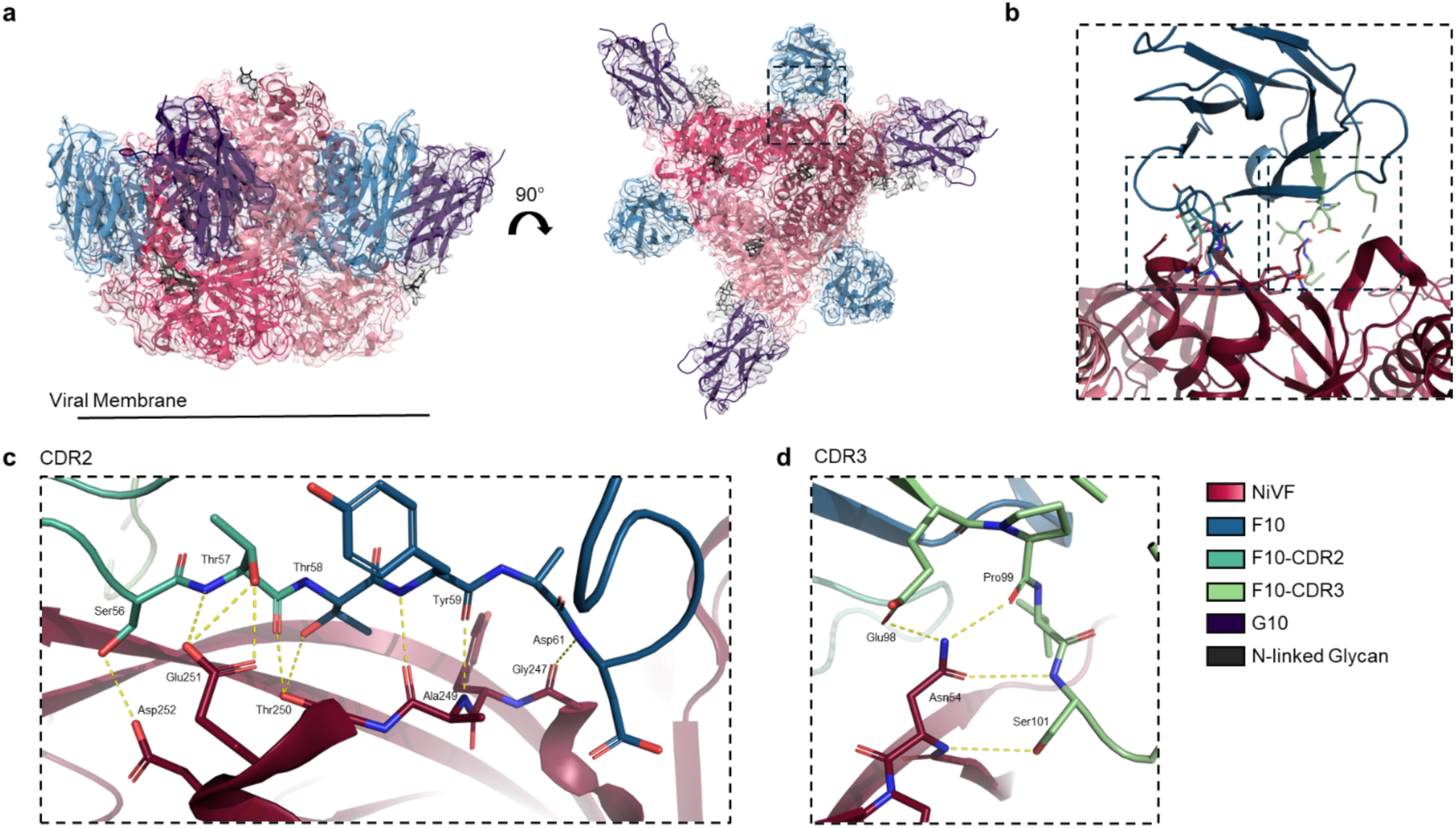
Epitope of F10 Nanobody. (a) Model fitting of NiVF, F10 and G10 into the Cryo-EM SPA volume, NiVF in red, F10 in blue, G10 in lilac and glycans in black. (b) Zoomed in view of the NiVF-F10 epitope. (c) Close up view of the CDR2 of F10 with interactions along a beta strand of NiVF. (d) Close up view of the CDR3 of F10 forming a H-bond network with NiVF. NiVF is in shades of pink, F10 is blue and the CRDs are in shades of green. Side chains of interacting residues are shown as sticks with hydrogen bonds labelled with dashes.

The D2 nanobody paratope covers an area of 720 Å^2^ of the NiVF head domain and comprises interactions involving all three complementarity-determining region (CDR) variable loops, with the majority contributed from the CDR2 and CDR3 loops. There is an extensive hydrogen bonding (H-bond) network from the backbone and sidechains of CDR2 of D2 with the sidechains of an alpha helix of NiVF (residues 73-85) (Fig. 4c,d). In particular, the sidechain of D2 Gln57 forms a H-bond with the sidechain of NiVF Glu77. The backbone of D2 Gln57 and D2 Arg58 forms H-bonds with the sidechain of NiVF Asn78 and the backbone of D2 Ala54 forms a H-bond with NiVF Thr81. The sidechain of D2 Arg56 is anti-parallel with NiVF Arg244 from the neighbouring protomer forming pi-pi stabilising interactions. The backbone of the D2 CDR3 residues Val101 and Met103 form H-bonds with the backbone of NiVF Gly247 and Ala249 respectively. The Phe102 residue of D2 fits into a hydrophobic pocket and forms pi-cation and pi-pi interactions with NiVF Lys55 and Phe282 respectively. D2 Asp105 forms a salt bridge with NiVF Lys55.

The F10 nanobody paratope covers an area of 903 Å^2^ of the NiVF head domain, overlapping with the D2 epitope, and shifted slightly towards the membrane. The interacting residues are mainly from CDR2 and framework region (FR) 3, forming a series of H-bonds between F10 Ser56 and F10 Asp 61 with NiVF Gly247 and NiVF Asp252 (Fig. 5c,d). The CDR3 of F10 forms 4 H-bonds with the sidechain and amide of NiVF Asn45.

The D2 nanobody epitope overlaps, but is more apical, with the epitope of the published nanobody DS90^30^. The F10 nanobody binds with a similar modality to the DS90 nanobody, with a H-bond network along the CRD2 and FR3. The CDR3s of F10 and DS90 bind in the same pocket of NiVF, forming H-bonds with different residues (Extended Data Fig. 9a-c).

### Conservation of the D2 and F10 epitopes in Henipahviruses

To assess whether the neutralising nanobodies D2 and F10 selected on the Malaysian variant of NiV would bind to Bangladesh NiV and other related henipaviruses, the F protein sequences of these viruses were aligned and the epitopes of the nanobodies mapped onto the alignment (Extended Data Fig. 10). The binding of D2 to NiVF involves 6 H-bonds, a salt bridge, a pi-cation and a pi-pi interaction and there is a total of 26 NiVF residues in the binding pocket. Comparing these residues across the two NiV strains and HeV shows that all are conserved between the NiV Malaysian and Bangladesh strains and there is only one conservative amino acid difference between NiV Malaysian and HeV, (Thr81 to Ser81), which forms a H-bond with Ala54 in CDR2 of D2. The epitope recognised by F10 includes 13 H-bonds across a binding interface comprising 28 residues. These residues are also conserved between NiV strains and HeV. There are several amino acid residues in the D2 and F10 epitopes that vary in other more distantly related henipaviruses such as Langya, Cedar, Yunnan Bat, Salt Gully and Ghana (Extended Data Fig. 10). From this analysis we conclude that the two neutralising nanobodies would bind to and neutralise both NiV strains and HeV.

## Discussion

NiV continues to be a threat with yearly outbreaks in Bangladesh and a recent outbreak in India, which is further aggravated by limited available therapeutics, prophylactics or point of care diagnostics^3,37^. Major efforts have focused on developing vaccines and therapeutics involving the two glycoproteins required for viral-host membrane fusion^23,26,27,38^. Of these, the F protein is more conserved between NiV strains and other henipaviruses than the G protein and therefore may be better suited as a target for cross-reactive antibody or nanobody therapeutics^6^.

In this study, we focused on the role of NiVF in the viral entry mechanisms, using cryo-electron tomography of NiV pseudoviruses and NiV VLPs to investigate the distribution of the NiVF on the surface membrane. Using sub-tomogram averaging, we determined the first structure of full-length NiVF, embedded in the viral membrane, and showed that NiVF dimer-of-trimers are present on the surface of NiV pseudoviruses and NiV VLPs. These appear to be part of a highly stochastic assembly of interacting NiVF-NiVF, not observed using purified NiVF ectodomain^21^, suggesting NiVF self-association is relatively weak and therefore transient. If this interaction was too strong, NiVF would be over-stabilised, held in the dimer-of-trimers, and never progress from the pre-fusion to the intermediate confirmation. The function of the NiVF dimer-of-trimers is still unknown, but it could stabilise NiVF in a pre-fusion conformation, transmit information regarding NiVG-ephrinB2 binding or help aggregate NiVF on the cellular membrane prior to budding. It could also help separate NiVF and NiVG into different areas of the virus.

A further possible function of NiVF assemblies is the sequestration of mature from immature NiVF; cathepsin-L cleavage between NiVF Arg109 and Leu110 during maturation would release the upstream fusion peptide, moving it apically, and towards the host membrane. The downstream residues would move basally and rest within the NiVF interface. This could mean mature NiVF preferentially form assemblies over immature, un-cleaved NiVF, regulating the endocytosis of isolated, immature NiVF and maintaining the mature NiVF on the host plasma membrane.

It is known that the binding of NiVG to EphrinB2 induces the conformational change of NiVF from the pre-fusion to the intermediate state, but the mechanism behind this is still unclear, with the two competing models, the *clamp* and the *provocateur,* differing as to whether NiVG and NiVF interact before or after receptor binding. The presence of NiVF assemblies on the surface of VLPs (and the lack of NiVG density) indicates limited to no binding of NiVF and NiVG on the surface of the VLPs prior to receptor binding, in keeping with the *provocateur* model. Evidence of these assemblies means we can expand on the current *provocateur* model adding a potential NiVF pre-fusion self-stabilisation step prior to receptor binding (Fig. 6c). However, further mechanistic studies are required to fully understand the relationship between NiVG-EphrinB2 binding and activation of pre-fusion NiVF. This process in the viral entry mechanism would also be an interesting target for future antibodies, nanobodies or organic molecules.

**Figure 6:**
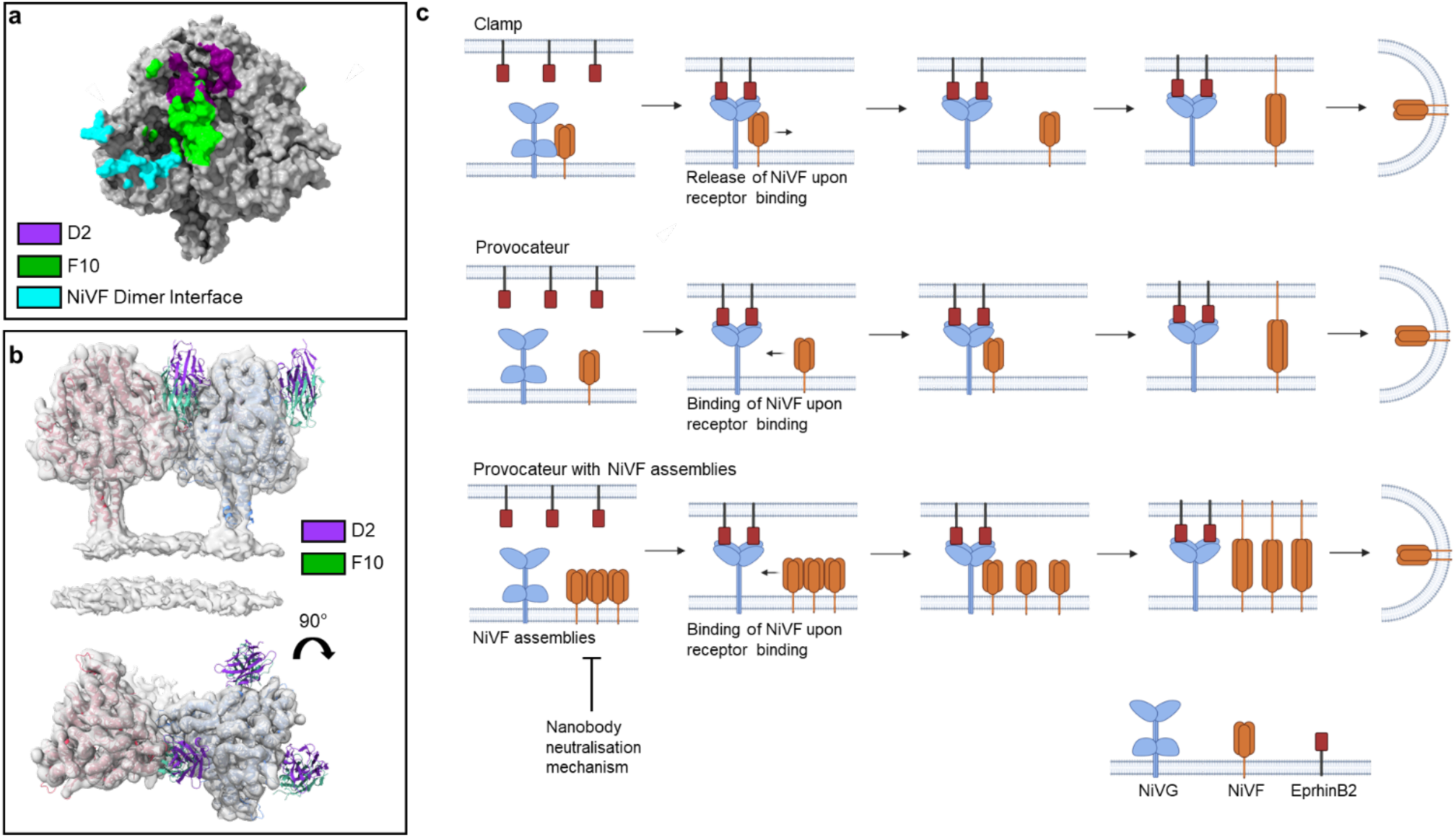
Neutralising Nanobodies Block NiVF-Interface. (a) A comparison of the D2 (purple), F10 (green) and the NiVF dimer-of-trimers epitopes (cyan). (b) NiVF dimer-of-trimer volume and atomic model with nanobodies superimposed from SPA atomic model. (c) A diagram with the NiV viral entry mechanisms: *clamp, provocateur* and *provocateur with NiVF assemblies.* Nanobody inhibition of NiVF assemblies is depicted to inhibit the formation of NiVF assemblies. NiVG, NiVF and EphrinB2 are depicted in blue, orange and red respectively.

We generated nanobodies against NiVF that exhibit high pseudovirus neutralising potency and high fusogenic-inhibition abilities. The nanobodies bound to two distinct epitopes on the NiVF ectodomain, however only one epitope was present on the wild-type NiVF. The neutralising nanobodies, D2 and F10, bind a quaternary crypt that has been found previously^30^, suggesting it is the most immunogenic site for nanobodies. The neutralisations of D2-Fc and F10-Fc were comparable with the other NiVF nanobody, DS90, and an anti-NiVG antibody: m102.4 kappa^30,31,39^. Comparison with published antibodies suggests D2 and F10 bind a similar location to the class of lateral antibody binders (Extended Data Fig. 9d). IC50 values of NiV VLP fusion inhibition and NiV HIV-1 pseudovirus neutralisation were comparable, suggesting the mechanism of action to be the same and therefore independent of NiVM or HIV Gag-Pol.

The major epitope of the neutralising nanobodies is close to the NiVF dimer interface observed by cryo-ET and therefore nanobody binding, in particular F10, would sterically prevent the formation of the dimer (Fig. 6a,b). NiVF is densely packed on the VLP, and it is also possible that the bivalent D2-Fc and F10-Fc constructs could cross-link NiVFs, thereby preventing the transition from pre-to post-fusion states. Therefore, the nanobodies may prevent fusion in the viral models by two mechanisms that are not mutually exclusive: namely, stabilisation of the pre-fusion conformation and the disruption of the dimer-of-trimers.

In summary, we have determined the high-resolution structure of full-length NiVF, integrated in the bilayer membrane of a VLP. We have provided evidence that NiVF forms assemblies on the VLP surface involving the association of dimers of trimers. The function significance of this is unclear but we speculate that these assemblies are required for viral entry. We have also developed highly-potent neutralising nanobodies that inhibit the fusion of NiV VLPs and pseudovirions, that are predicted to block the formation of the NiVF dimer-of-trimers, and with strong evidence that they have the potential as future therapeutics, prophylactics or point of care diagnostics.

## Methods

### NiV Pseudovirus Production

Neutralisation capabilities of nanobody constructs were determined using a NiV HIV-1 luciferase reporter pseudoviruses for which plasmids were kindly provided by Dalan Bailey from The Pirbright Institute. To generate NiV pseudoviruses, HEK293T cells were plated at 0.5 x 10^6^ cells/ml in DMEM 5% FBS medium one day before transfection. Cells were transfected by mixing plasmids pcDNA3.1(+) NiVF, pcDNA3.1(+) NiVG, p8.91 and pCSFLW at a ratio of 1:1:1:1.5 in OPTIMEM media. PEI was mixed with the DNA at a ratio of 3:1 and incubated for 10 mins at room temperature, before being added dropwise to the cells. 36 hours post-transfection, the media containing pseudoviruses was removed, centrifuged for 5 mins at 500 x g to remove cellular debris and filtered through a 0.2 μm filter.

To determine pseudovirus infectivity, one day before transduction, HEK293T cells were plated at 0.2 x 10^5^ cells/ml in DMEM 5% FBS medium in a Nunc MicroWell 96-well plate. On the day of transduction, pseudoviruses were added to the cells at varying dilutions in DMEM. This mixture was then transferred onto the cells and incubated for a further 48 hours. To measure luciferase expression, the media was discarded and 50 μl of 1:1 Bright-Glo™ (Promega):DPBS was added to the cells, and luminescence was measured after 5 mins by a CLARIOStar Plus (BMG Labtech).

### NiV VLP Production and Fusion Ability

A NiV VLP model was established to investigate NiV envelope protein distribution and interactions. The gene for NiVM fused to beta-lactamase (M-Blam), kindly provided by Stuart Neil, King’s College London, was codon-optimised and cloned into pcDNA3.1(+) vector using Infusion® cloning. Plasmids containing NiV M-Blam, NiVF and NiVG genes were transiently transfected into HEK293T cells at a 24:3:1 ratio using 5:1 DNA:PEI ratio. VLP-containing supernatant was harvested 48 hours post-transfection and clarified by centrifugation at 500 x g for 5 mins. VLPs were concentrated and purified via one or two rounds of 8-16 hours, 4000 x g centrifugations through a 10% sucrose cushion. The VLP pellet was resuspended in DPBS.

VLP fusogenic activity was tested by intracellular beta-lactamase activity via cleavage of a FRET-couple fluorophore pair, CCF2-AM. Briefly, wells were coated in polylysine at 0.1 mg/ml for 5 mins and HEK293T cells were seeded at 3 x 10^5^ cells/ml. The next morning, the media was replaced with concentrated VLPs diluted in DMEM and spinoculated by centrifugation at 2100 x g for 30 mins at 4°C. The cells were incubated for 90 minutes at 37°C, after which the media containing unfused VLPs was removed, and the cells were washed with DPBS. A solution of CCF2-AM and Probenecid, an organic anion transporter inhibitor, were added to the cells and incubated for 2 hours in the dark at room temperature, following manufactures instructions. After which the cells were washed with DPBS, detached from the well using Trypsin (0.05%), inactivated with 5% FBS DPBS and fixed using 4% PFA DPBS. Cells were counted onto a CytoFlex LX (Beckman Coulter) and fluorescence was measured using the FITC and Pacific Blue presets, with a minimum of 10,000 events per well. Analysis was carried out using FlowJo software (v. 10.10.0). Fusion was determined by the ratio of PacificBlue to FITC signal, an indicator of CCF2-AM cleavage and beta-lactamase activity (Supplementary Fig. 3).

### Cryo-Electron Tomography of NiV Pseudovirus and VLP

Quantifoil R1.2/1.3 200 Au grids were glow discharged for 30 seconds at 30 mA and 3 ul of purified NiV pseudovirus or VLP was applied to the grid. The grids were blotted for 3 seconds, at force +3, and flash-frozen in liquid ethane using a Vitrobot Mark IV (Thermo Fisher Scientific). Grids were manually screened for ice thickness, contamination, and sample density using a 200 kV Glacios 2 (Thermo Fisher Scientific) at Electron Biology Imaging Centre (eBiC), Diamond Light Source (Didcot, UK). Data were collected with Tomo5 software on a Titan Krios G4 cryo-transmission electron microscope (Thermo Fisher Scientific) operating at 300 kV with a Falcon 4i camera and a Selectris X energy filter detector, with a 10-eV slit, at the Rosalind Franklin Institute (Didcot, UK). For pseudoviruses, tilts were collected in a dose-symmetric scheme, ±55° in 2.5° increments. A dose of 170 e^-^/Å^2^ across the tilt series, at a magnification of 165000x (calibrated pixel size of 0.7584 Å). The defocus range used was from -3.0 to -5.0 in 1.0 increments. For VLPs: tilts were collected in a dose-symmetric scheme, ±60° in 3° increments. A dose of 140 e^-^/Å^2^ across the tilt series, at a magnification of 81000x (calibrated pixel size of 1.569 Å). The defocus range used was from -1.5 to -3.5 in 0.25 increments. Tilt-series were collected in EER format using multi-shot beam shift acquisition.

### Tilt Series Alignment and Tomogram Reconstruction

Tilt series were manually screened for poor quality images (grid bars, contamination or motion blur) before pre-processing for gain correction, CTF estimation, motion correction using a frame group size of 10 on Warp (v. 2.0.0)^40^. Tilt series were aligned using IMOD (v. 5.1.2)^41^ coarse alignment and fine alignment with 6 mm gold fiducials or using AreTomo (v. 2.0)^42^, and tomograms were reconstructed using Warp (v. 2.0.0). Tomograms were denoised using Isonet (v. 1.3)^43^ and VLP sizes were measured using IMOD (v. 5.1.2). VLPs were segmented using membrain-seg^44^.

### NiVF Particle Picking and Sub-tomogram Averaging on VLPs and Model fitting

Particles were picked manually using IMOD (v. 5.1.2) on tomograms at bin 8 or 16 and extracted using Warp (v. 2.0.0) at bin 4 or 8. Relion (v. 4.0.1)^45^ was used to align and classify sub-tomograms to remove junk and separate NiVF into different states. Further resolution improvements were made with iterative rounds of classification, refinement, re-centering and re-extraction of the NiVF dimer-of-trimers. NiVF dimer-of-trimers from VLPs were processed further with sub-tomogram CTF refinement, tilt-series refinement and temporal poses on M (v. 2.0.0)^40^.

For model fitting, a structure of signal-sequence cleaved NiVF (residues 27-546) was generated using AlphaFold2^33^ and roughly aligned into the volume map in ChimeraX (v1.7) before using an inbuilt fitinmap tool. The transmembrane domain was then removed using ChimeraX (v1.7) and the model (residues 27-486) roughly fit with one round of RealSpaceRefinement using Phenix (v. 1.20.1). Angles were measured using ImageJ (v. 1.8.0)^46^. Images were generated by ChimeraX (v1.7) with the Artiax plug-in (v0.6.0)^47^.

### Antigen Production

The pre-fusion stabilised ectodomain of NiVF (amino acids 26-488, with four single-point mutations (L104C, I114C, L172F and S191P), as previously described^34^, was ordered as a gene-block from IDT and inserted into pOPINTTGneo-GCNt vector using Infusion® cloning. The pOPINTTGneo-GCNt contains a mu-phosphatase leader sequence, a GCNt trimerization domain and C-terminal His6 tag and was generated in house^48^. NiVF plasmid was transfected into expi293™ with PEI and OPTIMEM, and the supernatant containing the desired protein was harvested 6 days post-transfection. NiVF was purified by tandem immobilized metal affinity chromatography (IMAC) and size exclusion chromatography (SEC) on an ÄKTAPure using a HiLoad™ 16/60 Superdex™200 or HiLoad 16/600 Superdex™200 column, using PBS pH 7.4 buffer. Fractions were assessed for purity by polyacrylamide gel electrophoresis (PAGE), pooled, concentrated, and flash frozen using liquid nitrogen.

### Llama immunisation and VHH library construction

Antibodies were raised in a llama as previously described^49^. NiVF protein (200 μg) was mixed with adjuvant (Gerbu LQ#3000) prior to intramuscular injections on days 0, 28, 56. Blood (150 ml) was collected on day 66. Immunization and llama handling were operated under the authority of the project licence PA1FB163A. cDNA, prepared form PBMCs, was amplified by two rounds of PCR and cloned into the Sfil sites of the phagemid vector pADL-23c (Antibody Design Laboratories, San Diego, CA, USA) generating a VHH library. Recombinant pADL-23c vectors were transformed into electro-competent *E. coli* TG1 cells (Agilent Technologies LDA UK), and the resulting TG1 library stock was infected with M13K07 helper phage to generate a library of VHH-presenting phages.

### VHH phage display and isolation

Two rounds of solid-phase bio-panning (50 and 5 nM of NiVF trimer) using 10 wells of a Nunc high-binding 96-well plate enriched pre-fusion NiVF specific VHHs which was confirmed with an Anti-llama polyclonal ELISA. Individual phagemid clones were picked from both pans, and the VHH-displaying phage were recovered by infection with M13K07 helper phage and binding to NiVF was detected by anti-M13 ELISA. The strongest binding clones were picked from the 50 and 5 nM pans, amplified by PCR, the DNA sequenced, and grouped by CDR3 sequence identity using the IMGT/V-QUEST server. Small scale expression of individual VHH clones was achieved by 1 mM IPTG induction in TG1 cells and growing overnight at 28°C. The crude VHH-containing periplasm was extracted using polymyxin and tested for NiVF binding using an anti-VHH ELISA. VHH clones were amplified using PCR from the phagemid vector and inserted into the pOPINVHH, pOPINVHH-cys or pOPINTTGneo-Fc vectors^50^, for monomer, monomer with a C-terminal Cys or recombinant-Fc expression respectively. pOPINVHH and pOPINVHH-cys contain an OmpA leader sequence and C-terminal His6 tag^51^. pOPINTTGneo-Fc contains a mu-phosphatase leader sequence, a huIgG1 Fc and C-terminal His6 tag^48^

### Nanobody and Recombinant Nanobody-Fc Production

VHH plasmids were transformed into the Wk6 *E. coli* strain and protein expression was induced by 1 mM IPTG and grown overnight at 28°C. Periplasm contents were extracted using osmotic shock and VHH proteins were purified by tandem IMAC and SEC on an ÄKTAPure using a HiLoad™ 16/60 Superdex™75 or Superdex™75 10/300 GL column, using PBS pH 7.4 buffer.

Recombinant Nanobody-Fc plasmids were transfected into expi29™ using PEI and OPTIMEM, and the supernatant containing the excreted desired protein was harvested 6 days post-transfection. Nb-Fcs were purified by Ni gravity columns, eluted using 300 mM imidazole and desalted using Zeba™ Desalting Spin Column (Thermo Fisher Scientific).

### ELISAs

High-binding 394-well plates were coated with 50 nM of NiVF ectodomain or purified NiV VLP (at a dilution previously determined) overnight and blocked with milk before incubation with Nb-Fc. An Anti-Fc antibody conjugated to horse radish peroxidase was used as the secondary antibody. Binding was determined by fluorescent readout of activated QuantaBlu™ (Thermoscientific); graphs were plotted and EC50 values were calculated using GraphPad Prism (v. 10.4.1).

### NiV VLP Fusion Inhibition Assays

A NiV VLP fusion assay was used to determine fusion inhibition of nanobodies. Fusion inhibition was determined by incubation of the VLPs with the purified protein for 90 mins prior to addition to the HEK293T cells. Inhibition was normalised with a CCF2-AM-only and a CCF2-AM and VLP-only sample and subsequent analysis was performed using GraphPad Prism (v. 10.4.1).

### NiV Pseudovirus neutralisation assays

To determine neutralisation capabilities, the same protocol for determining pseudovirus infectivity was followed. However, on the day of transduction, pseudovirus was incubated with nanobody/antibody constructs in DMEM for one hour at 37°C in 5% CO_2_. This mixture was then transferred onto the cells and incubated for a further 48 hours. Luciferase signal was then measured similarly to before, and the results were normalised with no-pseudovirus and pseudovirus-only controls.

### Biolayer Interferometry of NiVF Nanobodies

Biolayer interferometry was used to measure the binding constants of nanobodies to biotin-tagged NiVF immobilised on streptavidin sensors 0.1% BSA (w/v) in PBS, pH 7.4. Assays were performed in black Greiner 96-well plate with a volume of 200 μl per well using an Octet R8 (Sartorius) system and designed using Octet BLI Discovery (v12.2.2.20) software. Purified NiVF ectodomain was biotinylated using Biotinylation Kit / Biotin Conjugation Kit – Lighting Link (Ab20179), according to manufactures instructions. 50 nM NiVF was loaded onto streptavidin sensors, with cycling parameters of 30 s baseline, 600 s loading, 30 s wash, 600 s association and 600 s dissociation. Signal was background normalized, aligned to the Y axis using the average of the baseline step, inter-step correction with the dissociation step, and Savitzky–Golay filtering. For epitope binning, an association step (1200 s) of the first nanobody, was followed by a baseline (30 s) and an association of the second nanobody (300 s). Reduction of the *R*_max_ value by 50% indicates competitive binding. Nanobody-antigen binding was modelled using 1:1 binding, with *K*_d_, *k*_a_ and *k*_dis_ values calculated using Octet Analysis Studio (v12.2.2.26), n=3. All graphs were plotted in GraphPad Prism (v. 10.4.1).

### Fluorescent nanobody labelling of NiVF on cell membranes

Nanobodies (D2, F10 and G10) expressed in pOPINVHH-cys, were reduced using 5 mM TCEP for 30 mins and labelled using Atto594-maleimide (Atto-Tec) following manufacturer’s instructions. To express NiVF: 20,000 HEK293T cells were seeded in an Ibidi 8-well (Ibidi) 1 day prior to transfection. 200 ng of pcDNA3.1(+) NiVF DNA was mixed with 0.4 ul FuGene in OPTIMEM, incubated for 10 mins and added to cells dropwise. 48 hours post-transfection, Nb-Atto594 was added to the cells and incubated for 15 mins. Cells were then imaged using a Leica SP8 Confocal Microscope (Leica) with a 60× oil immersion objective and HeNe 594 nm laser, and images were processed using ImageJ (v. 1.8.0).

### Fluorescence Lifetime Cross-Correlation Spectroscopy of GFP-labelled Pseudovirus with Atto594-laballed Nanobodies

Fluorescence lifetime cross-correlation Spectroscopy (FLCCS) was performed using the MicroTime200 (Picoquant) time resolved fluorescence microscope. Serial dilutions of purified pseudoviruses labelled with Gag-eGFP were incubated with 5 nM of single domain antibodies labelled with Atto594 in a final volume of 30 uL in PBS, using low-binding tubes during 1 h at 37 °C. The sample was transferred to a glass chambered coverslip immediately prior to FLCCS acquisition (Cat.No:81506, Ibidi). The incubator chamber (DigitalPixel) was preheated to 37°C and the sample was excited using pulsed 485 nm (for HIV-1 virions labelled with Gag-eGFP) and 595 nm (for single domain antibodies labelled with Atto594) diode lasers (LDH series Picoquant) with a repetition rate of 20 MHz. The laser beam was coupled to an Olympus IX73 inverted microscope and focused onto the sample by a 63× 1.2 NA water immersion objective lens (Olympus UPlanXApo). Emission passed through a quad-dichroic mirror specifically designed to reflect four installed laser excitation lines (at 440, 485, 594, and 635 nm, Chroma) and a 100 μm pinhole (ThorLabs). The remaining emission was separated at a 560LP dichroic (Chroma). The longer-wavelength emission was then passed through 600LP and 690/70BP (Chroma) before arriving at a SPAD (Picoquant). The shorter-wavelength emission was passed through a 525/50B filter (Chroma) before a PMA hybrid detector (Picoquant). Time correlated single photon counting (TCSPC) was performed using the Multiharp 150 (Picoquant). The autocorrelation and cross-correlation curves for single viruses, single domain antibodies and single viruses engaged with single domain nanobodies were recovered and analysed with SymPhoTime software (Picoquant) employing a 2-diffusion Triplet Extended (3D) model and lifetime filters for FLCCS^36,52^.

### Cryo-electron Microscopy of NiVF-Nb complexes

Purified NiVF at 0.2 mg/ml was mixed with a 1:3.6 molar ratio with a nanobody in PBS and incubated on ice for 30 minutes. 3 ul was applied to a glow-discharged Quantifoil R 1.2/1.3 on 200 Au mesh grid. Grids were blotted using a Vitrobot Mark IV (Thermo Fisher Scientific) for 3 seconds and with +3 or - 10 blot force, and plunge frozen in liquid ethane. Grids were screened for ice thickness, contamination, and particle density using a 200 kV Glacios 2 (Thermo Fisher Scientific) at the UK Electron Biology Imaging Centre (eBiC), Diamond Light Source (Didcot, UK). Data were collected using EPU on a Titan Krios G4 cryo-transmission electron microscope (Thermo Fisher Scientific) operating at 300 keV with a Falcon 4i camera and a Selectris X energy filter detector with an electron dose of 50 e^-^/Å^2^ at a 0° or 30° tilt and a magnification of 165000x (calibrated pixel size of 0.7584 Å) at the Rosalind Franklin Institute (Didcot, UK). Micrographs were corrected for motion blur and CTF estimation using CryoSparc (v 4.7.1). Particles were initially picked using Cryosparc’s in-built blob-picking before switching to template picking after a 2D classification of a subset of particles. 3D volumes were generated using ab-initio and refined using iterative rounds of heterogenous refinement, homo-refinement, global and local CTF refinement. Rebalance orientation was also employed for the NiVF-F10-G10 complex due to preferential orientation.

A NiVF model (PDB: 7upd^25^) was roughly aligned into the volume map in ChimeraX (v. 1.7) before using an inbuilt fitinmap tool and manual repositioning of atoms in Coot (v. 0.9.8.95). Nanobody protein template structures were generated using an online Nanobody builder tool^53^ and manually rebuilt into the volume map using Coot (v. 0.9.8.95)^54^. Atomic models were combined in ChimeraX (v. 1.7) and iterative rounds of model building, and real space refinement were carried out using Coot and Phenix (v. 1.20.1). Images were created using ChimeraX (v. 1.7) or Pymol (v. 3.1.6.1).

### Statistics and Reproducibility

Statistical analysis was performed using GraphPad Prism (v. 10.4.1). IC50 and EC50 values were calculated using a sigmoidal curve analysis with four parameters, with background subtraction and normalisation where appropriate. Three repeats were carried out, over two experimental days. FLCCS Nb Cross-correlation and slow-diffusion coefficient was plotted with a sigmoidal curve analysis with four parameters. Frequency of Nb detection events was plotted with a damped model using the custom equation, *𝑌 = 𝑌_0_ + 𝐴(𝑒^-k^*^1x^ *– 𝑒*^-k2x^).

## Data Availability

Anti-NiVF VHH DNA Sequences have been deposited in GenBank with the accession numbers: PZ374715-PZ374726. Cryogenic electron microscopy maps and models have been deposited in the Electron Microscopy Data Bank (EMDB) and the Protein Data Bank (PDB) as follows: NiVF+D2 (EMD-57949, 30QG); NiVF+F10+G10 (EMD-57950, 30QH). Pseudovirus NiVF Dimer Cryogenic electron tomography map has been deposited in the EMDB as follows: NiVF Dimer (EMD-57952). VLP NiVF Dimer Cryogenic electron tomography map and model have been deposited in the EMDB and the PDB as follows: NiVF Dimer (EMD-57951, 30QI).

## Acknowledgement

This work was supported by the Rosalind Franklin Institute, funding delivery partner EPSRC, an EPSCR Studentship Grant (EP/W52430X/1) and Wellcome Trust (Grant No. 223733/Z/21/Z). S.P.-P. acknowledges funding from the European Research Council (Grant No. ERC-2019-CoG-863869 FUSION) and The Chan Zuckerberg Initiative “Multi-color single molecule tracking with lifetime imaging” (Grant No. 2023-321188). We thank our colleagues at the Rosalind Franklin Institute, Liang Wu for help with initial collection of tomograms, Casper Berger, and Helena Watson for assistance with Cryo-ET data processing, John. Clarke for advice with cryo-EM data processing, Lauren Eyssen for providing the phage display library and Hannah Campaigne for assistance in seroconversion assays and VHH panning. We also thank Stuart Neil (King’s College, London) for advice on VLPs and Gary Stephens, Barney Jones, Hong Lin (Reading University, UK) for expertise in llama immunization.

## Author contributions

Nanobodies were identified and all antigen-binding assays carried out by J.W.T, fluorescent correlation spectroscopy experiments were performed and analysed by I.C-A.. J.W.T. carried out neutralisation assays following methods developed and validated by L.J., Cryo-EM and cryo-ET was performed and analysed by J.W.T.. R.J.O., M.C., S.P-P and J.W.T. planned the project and interpreted the data. J.W.T, S.P-P and R.J.O wrote the paper with contributions from all authors.

## Extended Data Figure Legends

**Extended Data Figure 1:**
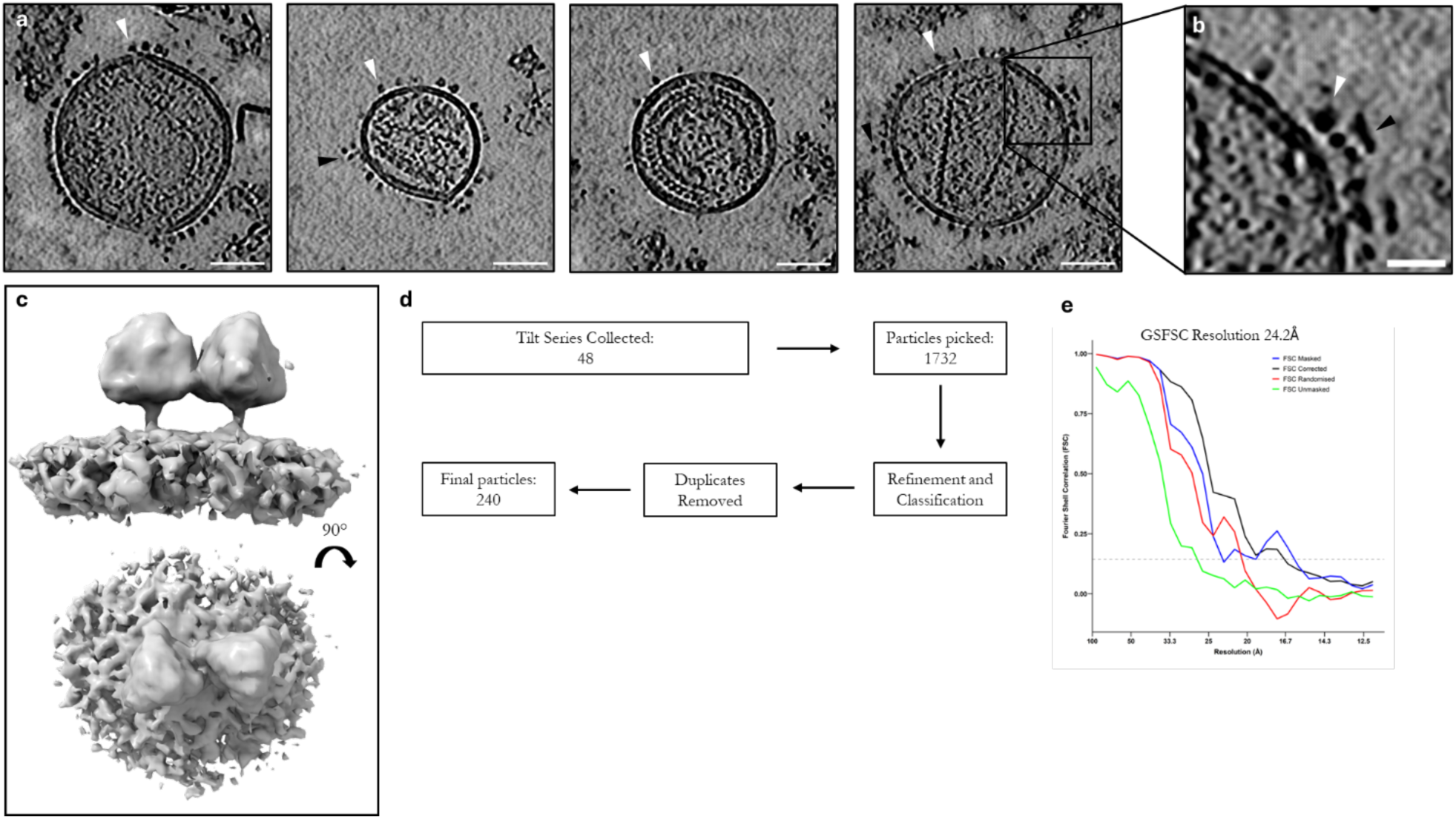
NiV Pseudovirus with NiVF Dimer-of-trimer. (a) Tomogram slices of NiV pseudoviruses. NiVF is depicted with a white arrow and NiVG with a black arrow. Scale bar is 50 nm. (b) Zoomed in image of NiV pseudovirus. Scale bar is 20 nm. (c) Cryo-ET of NiVF dimer-of-trimers seen on NiV pseudovirus. (d) Pipeline of sub-tomogram averaging. (e) GSFSC curve of NiVF dimer-of-trimers.

**Extended Data Figure 2:**
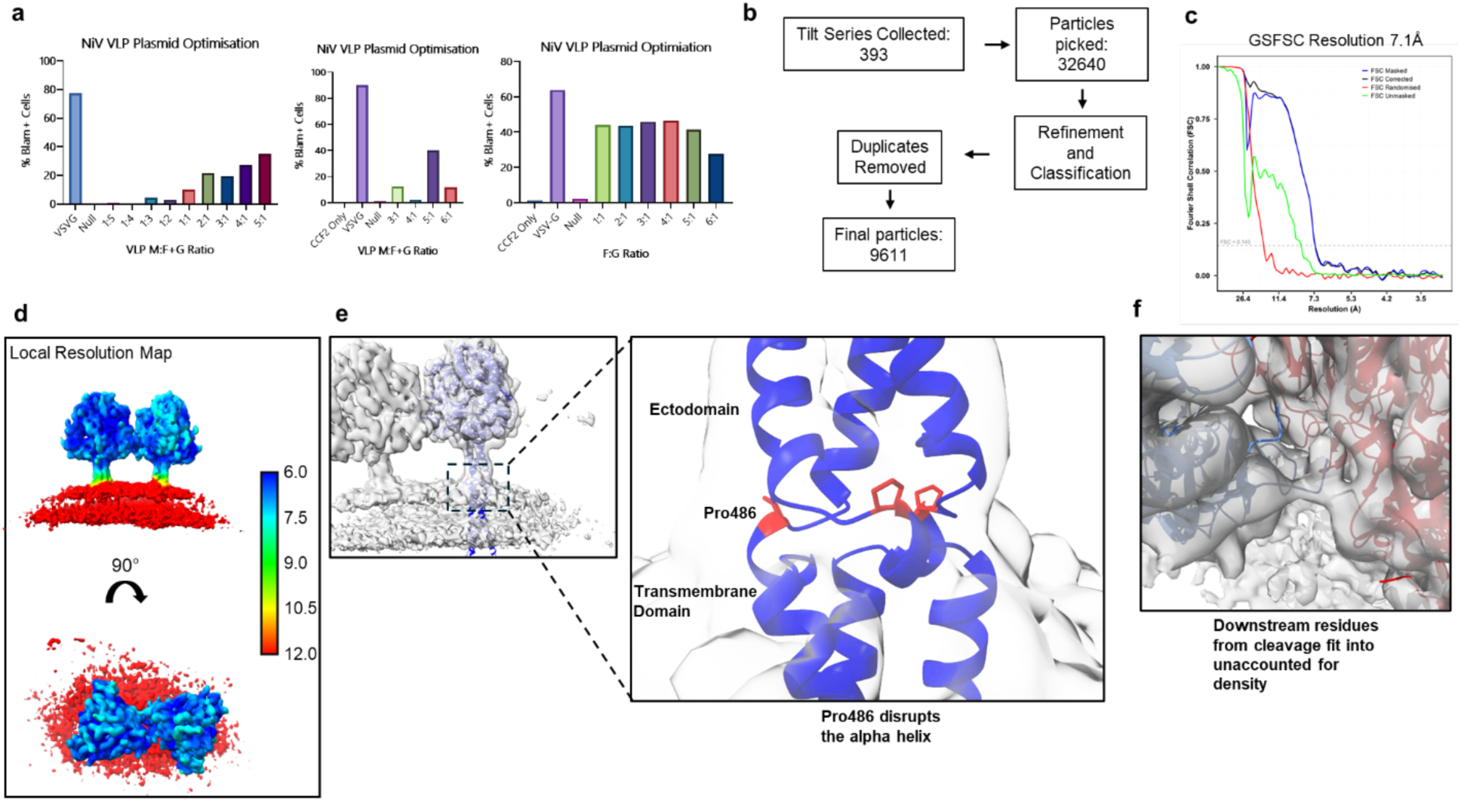
VLP Generation and Cryo-ET Analysis. (a) Optimisation of plasmid ratios of NiVF, NiVG, and NiVM based on maximising fusogenicity of NiV VLPs. (b) Processing pipeline including number of micrographs and particles. (c) Gold-standard Fourier shell correlation (GSFSC) after FSC-mask auto-tightening calculated by M (v. 2.0.0). The grey horizontal dashed line represents the 0.143 threshold^55^. (d) NiVF dimer-of-trimers local resolution map. Scale bar is in in Å. (e) NiVF Pro486 highlighted red in the full length NiVF AlphaFold model suggesting disruption of the alpha helices between the ectodomain and transmembrane domain. (f) Downstream residues from Cathepsin L cleavage site in unaccounted for density in NiVF dimer-of-trimer model.

**Extended Data Figure 3:**
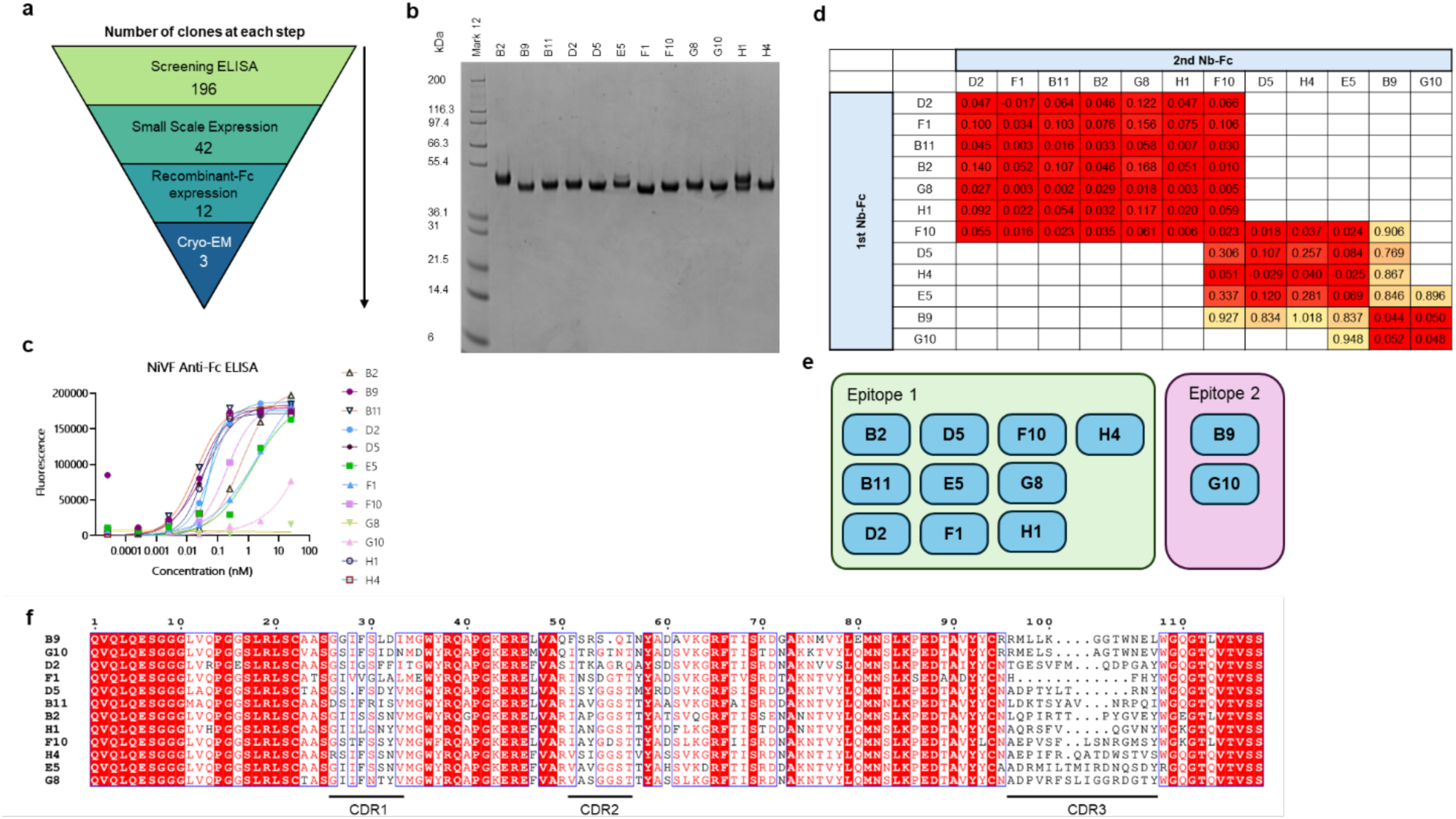
VHH isolation and biochemical characterisation. (a) The funnel explaining the reduction in clones throughout the VHH isolation workflow. (b) SDS-PAGE of 12 Nb-Fc clones after mammalian cell expi293T™ expression and Protein A purification. (c) NiVF Anti-Fc ELISA of the 12 Nb-Fc clones. (d-e) Epitope binning matrix obtained using bio-layer interferometry with corresponding binning map. (f) Summary of 12 nanobodies expressed recombinantly and their corresponding amino acid sequences with labelled CDRs. Image made using ESPirit^56^.

**Extended Data Figure 4:**
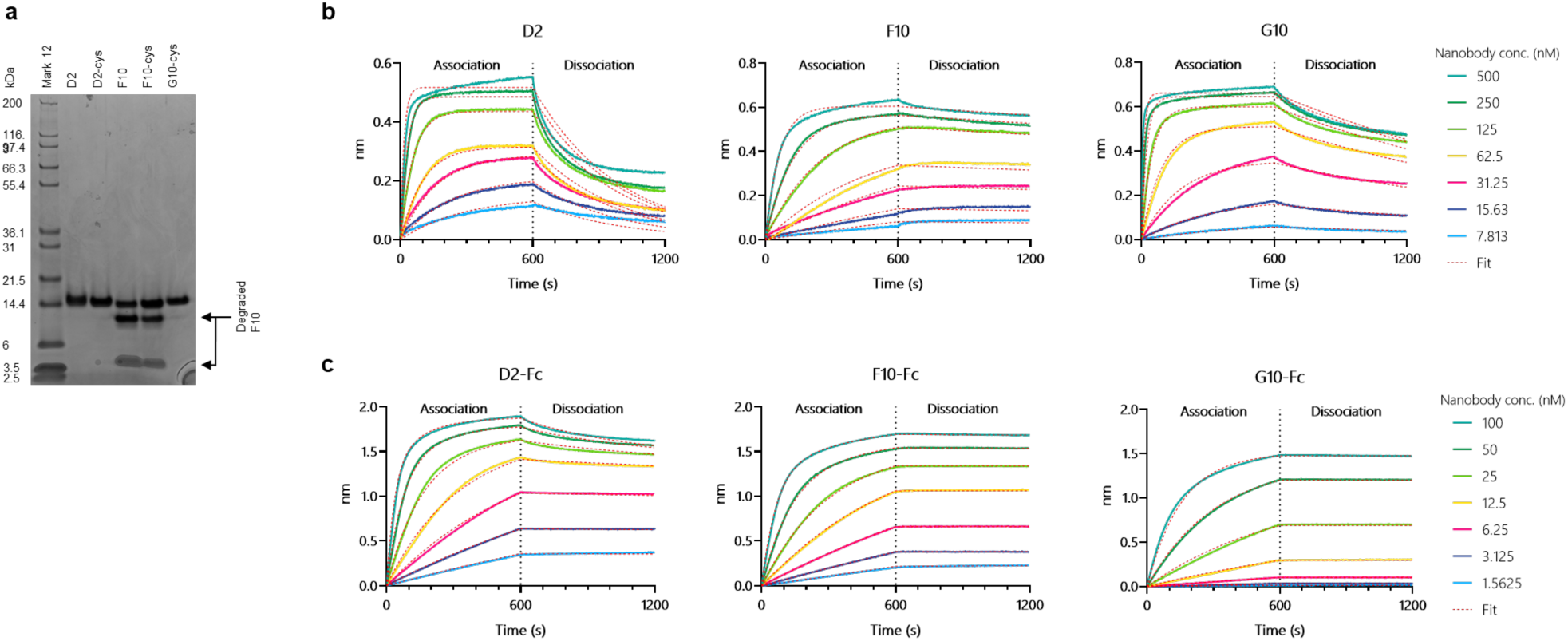
VHH characterisation and bio-layer interferometry Data. (a) SDS-Page of D2, F10 and G10 from bacterial *E. coli* Wk6 expression and tandem IMAC and SEC purification. Samples molecular weight was measured by time-of-flight mass spectroscopy: experimental molecular weights agreeing with calculated masses. D2 and G10 were purified as a single protein species, while F10 had a secondary contaminant, with the same molecular weight as a degraded F10 species. (b) Bio-layer interferometry of D2 and F10 single-domain nanobodies. Both D2 and F10 have low nanomolar to sub-nanomolar affinity calculated as a 1:1 binding; F10 has a stronger affinity to NiVF ectodomain than D2. (c) Bio-layer interferometry of D2, F10 and G10 with recombinant Fc domains. Recombinant Nb-Fc species have a greater affinity than nanobody monomers.

**Extended Data Figure 5:**
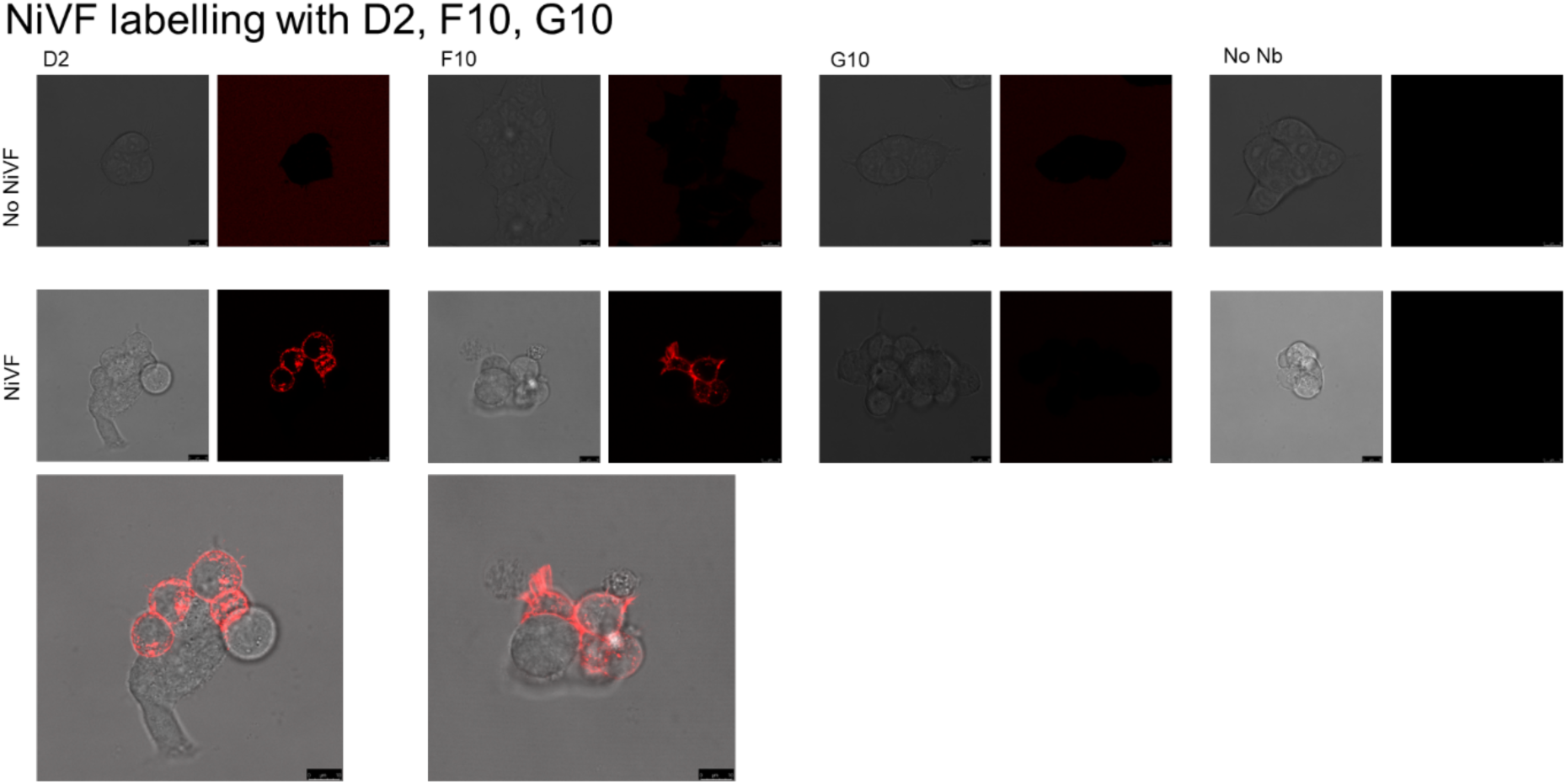
Cell labelling of NiVF using Atto594-labelled nanobodies. (a) Atto594 labelled NiVF nanobodies showing binding to wild-type NiVF expressed on the surface of HEK293T cells. D2 and F10 binding the wild-type NiVF, while G10 shows no binding to wild-type NiVF.

**Extended Data Figure 6:**
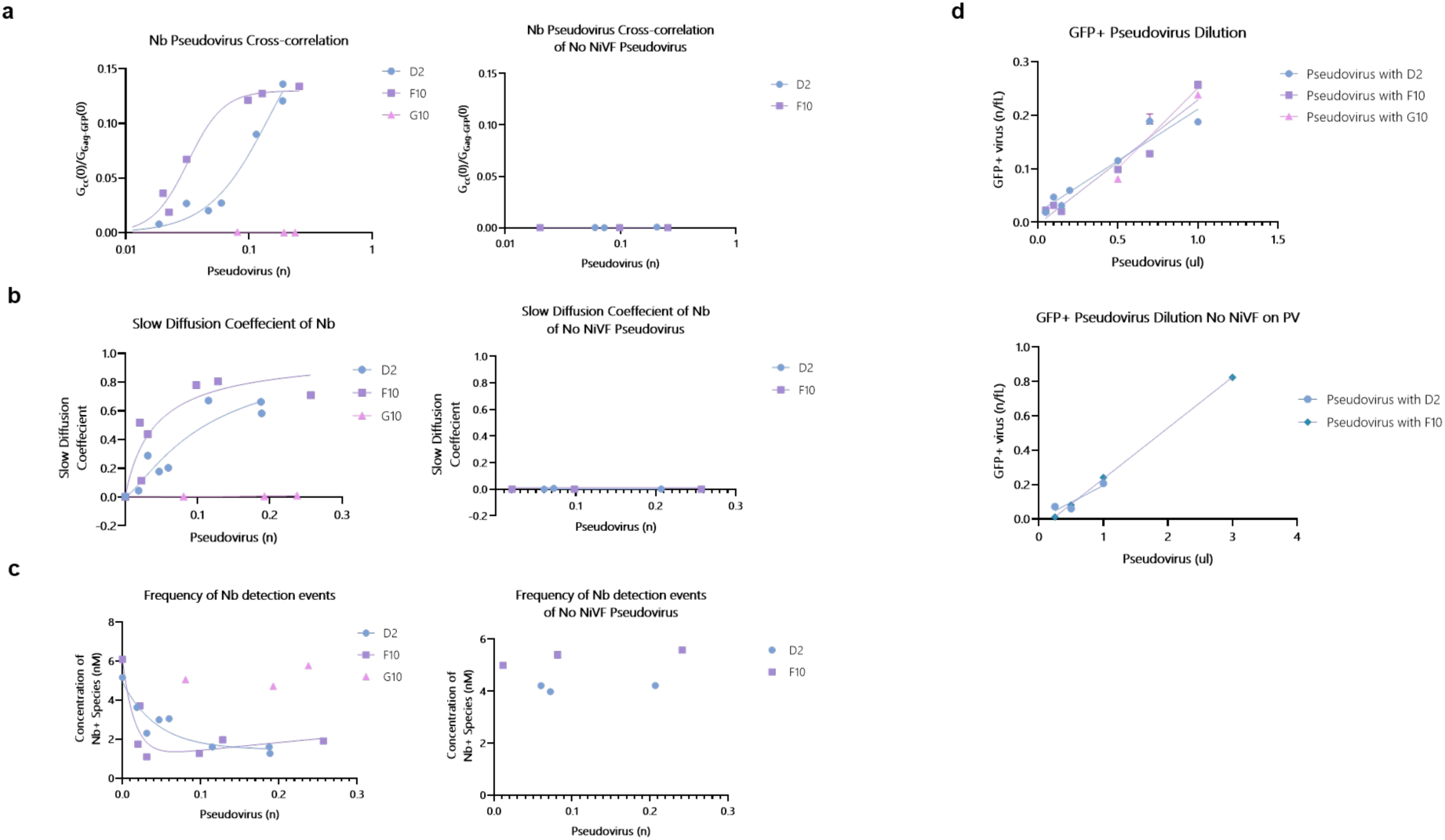
Fluorescent Cross-Correlation Spectroscopy Data and Controls. (a) Atto594-labelled nanobody cross correlation graphs. (b) Slow diffusion coefficient of nanobody shows D2 and F10 both binding pseudoviruses but no binding of G10. As pseudoviruses are added, nanobodies bind and diffuse at a slower rate. (c) Frequency of nanobody detection events against concentration of pseudoviruses. As pseudoviruses are added, the nanobodies are sequestered resulting in fewer detection events. (d) Dilution curve of pseudoviruses and no NiVF pseudoviruses control shows linear dilution curve. Controls with pseudoviruses expressed with no NiVF also shown.

**Extended Data Figure 7:**
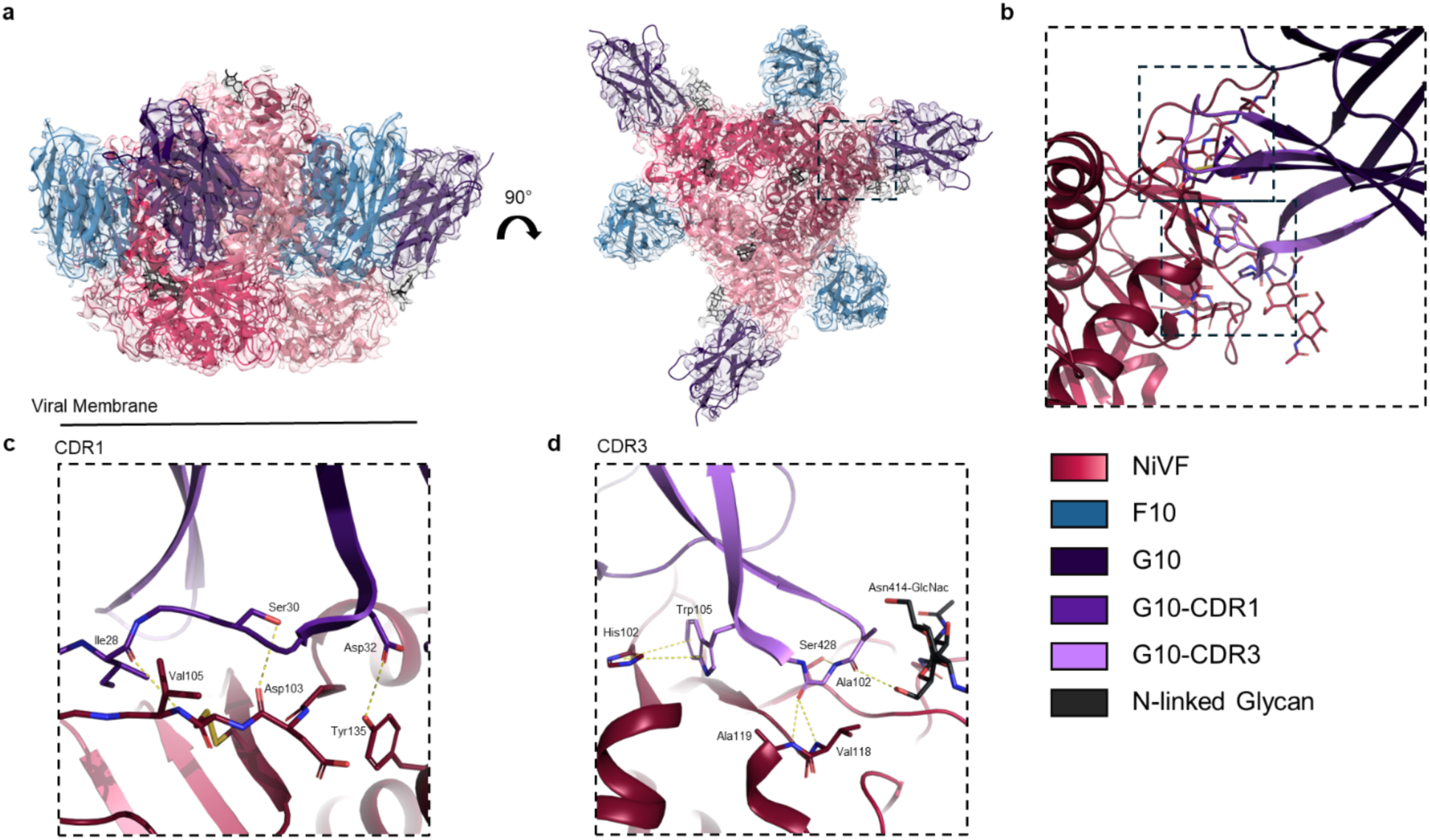
Epitope of G10 Nanobody. (a) Model fitting of NiVF, F10 and G10 into the Cryo-EM volume, NiVF in red, F10 in blue, G10 in lilac and glycans in black. (b) Zoomed in view of the NiVF-G10 epitope. (c) Close up view of the CDR1 of G10 interacting with residues either side of the cysteine mutation stabilising the flexible loop (Leu104Cys). (d) Close up view of the CDR3 of G10 forming a pi-stacking interaction and H-bonds with NIVF amino acid residues. Also shown is a H-bond from G10 to an N-linked Glycan on NiVF. Side chains of interacting residues are shown as sticks with hydrogen bonds labelled with dashes.

**Extended Data Figure 8:**
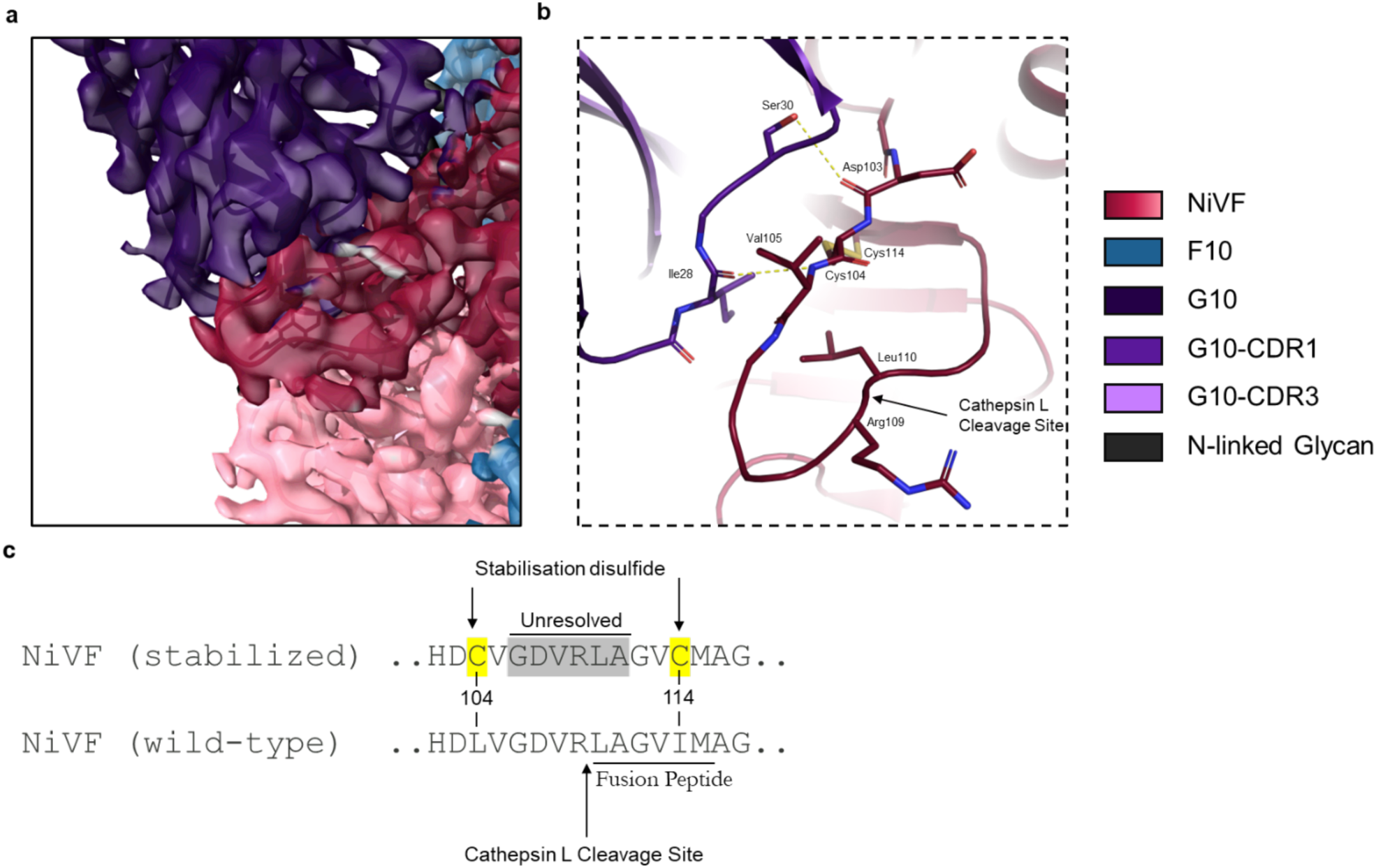
Cryo-EM Structure of NiVF:G10 stabilised loop. (a) Map volume of NiVF-G10 including the stabilised loop (red). (b) Atomic model of the stabilised loop with the disulfide bond (c) Amino acid sequence of the stabilised NiVF loop and the wild-type loop. Region often unresolved in cryo-EM SPA structures also highlighted.

**Extended Data Figure 9:**
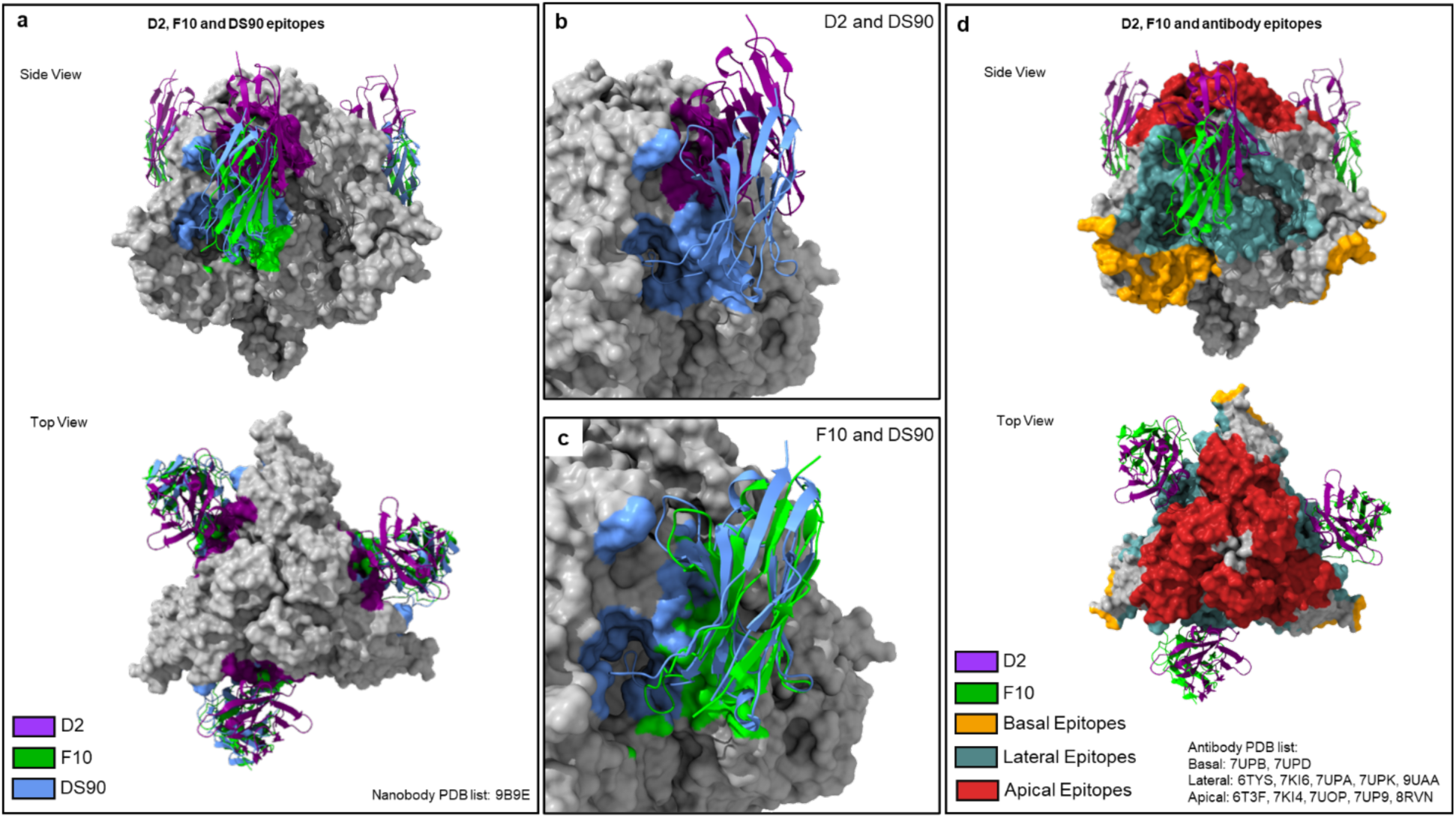
D2 and F10 epitope comparison. NiVF PDB superimposed with other NiVF antibody and nanobody structures (DS90)^30^. (a-c) D2, F10 and DS90 epitopes are highlighted in purple, lime and blue respectively. (d) Antibodies epitopes are highlighted based on basal, lateral or apical in red, cadet blue and orange respectively^22–27,30^.

**Extended Data Figure 10:**
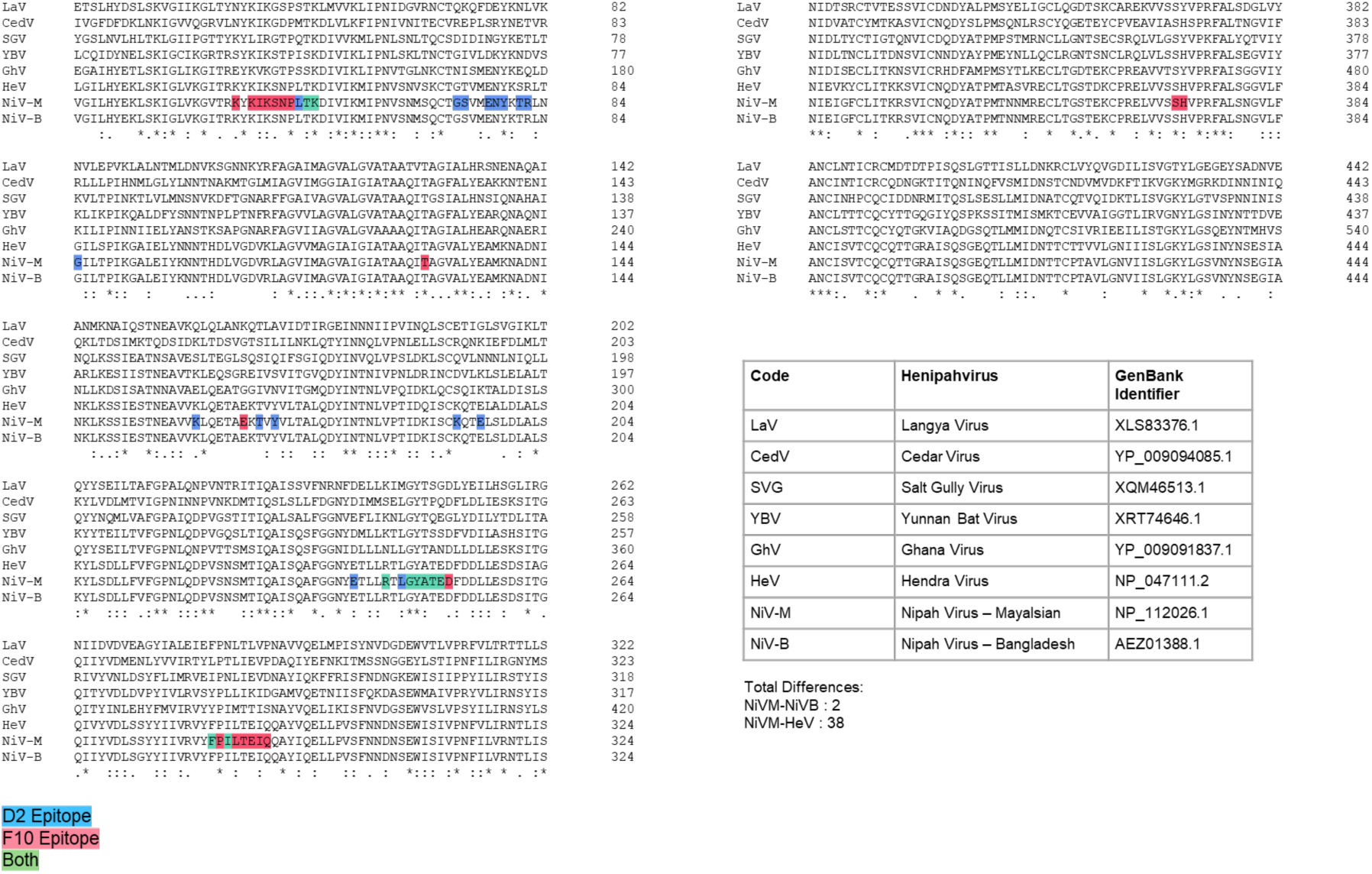
Multiple sequence alignment of D2 and F10 Epitopes on NiVF Ectodomain. (a) Multiple sequence alignment of amino acid sequences of the F proteins of common henipaviruses. Residues highlighted based on the D2, F10 epitopes or both, in blue, red and green respectively. Includes table of GenBank identifies for each henipavirus.

